# Coiled-coil heterodimers with increased stability for cellular regulation and sensing SARS-CoV-2 spike protein-mediated cell fusion

**DOI:** 10.1101/2020.12.10.419440

**Authors:** Tjaša Plaper, Jana Aupič, Petra Dekleva, Fabio Lapenta, Mateja Manček Keber, Roman Jerala, Mojca Benčina

## Abstract

Coiled-coil (CC) dimer-forming peptides are attractive designable modules for mediating protein association. Highly stable CCs are desired for biological activity regulation and assay. Here, we report the design and versatile applications of orthogonal CC dimer-forming peptides with a dissociation constant in the low nanomolar range. *In vitro* stability and specificity was confirmed in mammalian cells by enzyme reconstitution, transcriptional activation using a combination of DNA-binding and a transcriptional activation domain, and cellular-enzyme-activity regulation based on externally-added peptides. In addition to cellular regulation, coiled-coil-mediated reporter reconstitution was used for the detection of cell fusion mediated by the interaction between the spike protein of pandemic SARS-CoV2 and the ACE2 receptor. This assay can be used to investigate the mechanism and screen inhibition of viral spike protein-mediated fusion under the biosafety level 1conditions.

## Introduction

The coiled-coil (CC) motif, ubiquitous in natural proteins, is characterized by the relatively well-defined assembly and specificity rules and is highly attractive as a building block for designing and developing novel polypeptide scaffolds ^1^, functional proteins ^2–5^, and as a regulator of protein-protein interactions ^6–8^. CC protein origami (CCPO) cages ^9,10^, polymer–peptide hydrogels, nanoparticles, and peptide fibers ^11^ are fine demonstrations of the utility of CCs for engineering new polypeptide nanostructures. The specificity of CC pairs has been harnessed in order to develop logic circuits and controllers, as well as mammalian cell engineering ^4,12^, however different applications, require different CC stabilities ^13–18^.

Since the stabilities of several previously-designed four-heptad long CC pairs either had low orthogonality to other proteins (e.g., E/K pair ^13^) or were only moderately stable ^16,19^, we sought to design a set of highly stable orthogonal parallel heterodimeric CCs that could improve the sensitivity and increase the range of existing or prospective biological applications.

The COVID-19 pandemic caused by SARS-CoV-2 infection is currently among the most acute problems of humanity^20^, and investigation of its function and discovery of inhibitors command the attention of many scientists. The enveloped SARS-CoV-2 viruses require fusion of viral and cellular membranes in order to infect host cells^21^. The membrane fusion process is assisted by viral fusion proteins located at the viral envelope. After binding of viral fusion proteins to the receptor at the host cell membrane, a structural rearrangement of the viral fusion protein lowers the energetic barrier of the coalescence of the two biological membranes and the introduction of the viral genome into the cytosol occurs ^22^. Multiple methods are used for the evaluation of the viral entry and alternative steps of this process ^23^, from active virus to pseudotyped virus (a.k.a. pseudovirus) infection, which require BSL-2 or BSL-3 labs in order to prevent the danger of infection. Alternatively, cell-cell fusion assays can be used, based on the formation of syncytia between cells that express viral fusion (spike) proteins and the cellular receptors (ACE2) on other cells, offering a safer, virus-free alternative for viral entry studies and the development of antiviral drugs. Syncytium formation has been traditionally analyzed by slow and semi-quantifiable microscopy ^24–26^. Generation of a signal, which occurs only after the fusion between donor and acceptor cells ^27^, provides a good basis for this sensitive method to monitor syncytium formation. Therefore, split fluorescence or luminescence proteins linked to synthetic dimerization domains that form very stable dimers would be a prime choice as reporters.

In order to generate a set of CC heterodimers with the high thermodynamic stability necessary for new biological applications, we built on a previously published orthogonal NICP CC set ^17^ but increased the stability of the peptide pairs by reducing the number of asparagine (Asn, N) residues ^28,29^ and modifying amino-acid residues at the noncontact *b*, *c*, and *f* positions ^16^. CC pairs were able to reconstitute a split reporter enzyme, and we additionally demonstrate that their activity could be regulated externally by the addition of peptides to cells. Similar to the split reporter reconstitution, we also demonstrated reconstitution of a CC split transcription factor. Moreover, we sought to employ the designed CC pairs in monitoring membrane fusion based on the interaction between SARS-CoV-2 spike protein and ACE2 receptor interactions. This sensitive syncytium formation assay can be used to monitor the fusion process and screen for inhibitors in a safe and easy-to-employ way.

## Results

### *De novo* designed orthogonal heterodimeric CC pairs with high stability

Using previously established principles ^16,17^, we designed four heptad long CC dimer-forming peptides (named N5–N8) expected to form two orthogonal CC pairs in a parallel orientation with a high binding affinity (**Table 1**, N-variants; a CC N7:N8 structural model is depicted in **Fig S1A**). While Asparagine–Asparagine (Asn–Asn) pairing at the *a* positions is important for ensuring interaction specificity and peptide orientation ^13^, it contributes unfavorably to the affinity ^28,30^. Therefore, in order to increase the affinity of the NICP set, we maintained only a single Asn–Asn interaction per CC pair. The Asn (N) residue was placed at the *a* position in the third heptad for peptides N5 and N6 and in the second heptad for peptides N7 and N8 to ensure their mutual orthogonality. All other *a* positions were occupied by isoleucine (Ile, I), whereas leucine (Leu, L) was placed at the *d* position in all heptad repeats. Given that the residues at the surface-exposed positions do not contribute to the pairing interactions ^16^, alanine (Ala, A) residues were placed at the surface-exposed *b*, *c*, and *f* positions, in order to promote high helical propensity and, consequently, increase CC stability. In order to ensure orthogonality, lysine (Lys, K) and glutamic acid (Glu, E) were positioned at the *e* and *g* positions, obtaining peptide pairs with a unique electrostatic motif that should promote the formation of two CC heterodimers named N5:N6 and N7:N8—both in parallel orientations. The electrostatic motifs of these N-type peptides were kept similar to those of the previously published NICP P-type peptides (P5–P8) ^17^ (**Table 1**, **Table S1**). The interaction pattern of the designed set was evaluated by the scoring function ^31^, supporting the favorable formation of N5:N6 and N7:N8 CCs (**Fig S1B**).

**Table 1.**
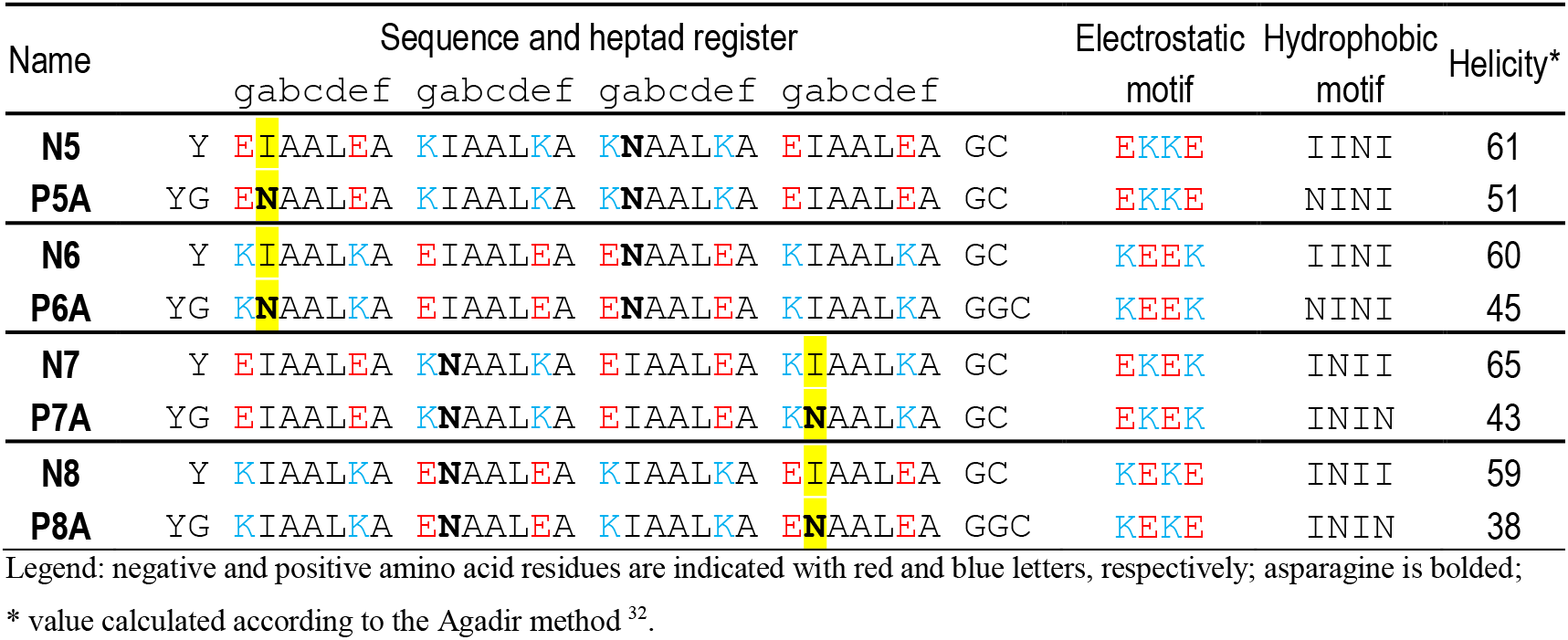
List of N- and PA-type parallel heterodimers.

Following peptide design, thermal stability and orthogonality of synthetic N-type peptides were analyzed. The secondary structure and stability of peptides were probed by circular dichroism (CD) spectroscopy. The designed peptide pairs, N5:N6 and N7:N8, were orthogonal to each other (**Fig 1E**), with the characteristic α-helical CD spectra, Tm above 70 °C (**Fig 1A**), and a dissociation constant determined by the ITC (K ^ITC^) in the low nanomolar range (1–15 nM; **Fig 1B**). In addition, size-exclusion chromatography coupled to static light scattering (SEC-MALS) analysis showed that the equimolar mixture of peptides preferentially formed heterodimers (**Fig S2B**); however, all four individual N-peptides were α-helical at 20 °C (**Fig S2A**). The melting curves of peptides were sigmoidal with a Tm midpoint between 37 °C and 55 °C (**Fig 1A**). Analysis by SEC-MALS confirmed the homodimeric formation of all peptides in solution (**Fig S2B**).

**Figure 1.**
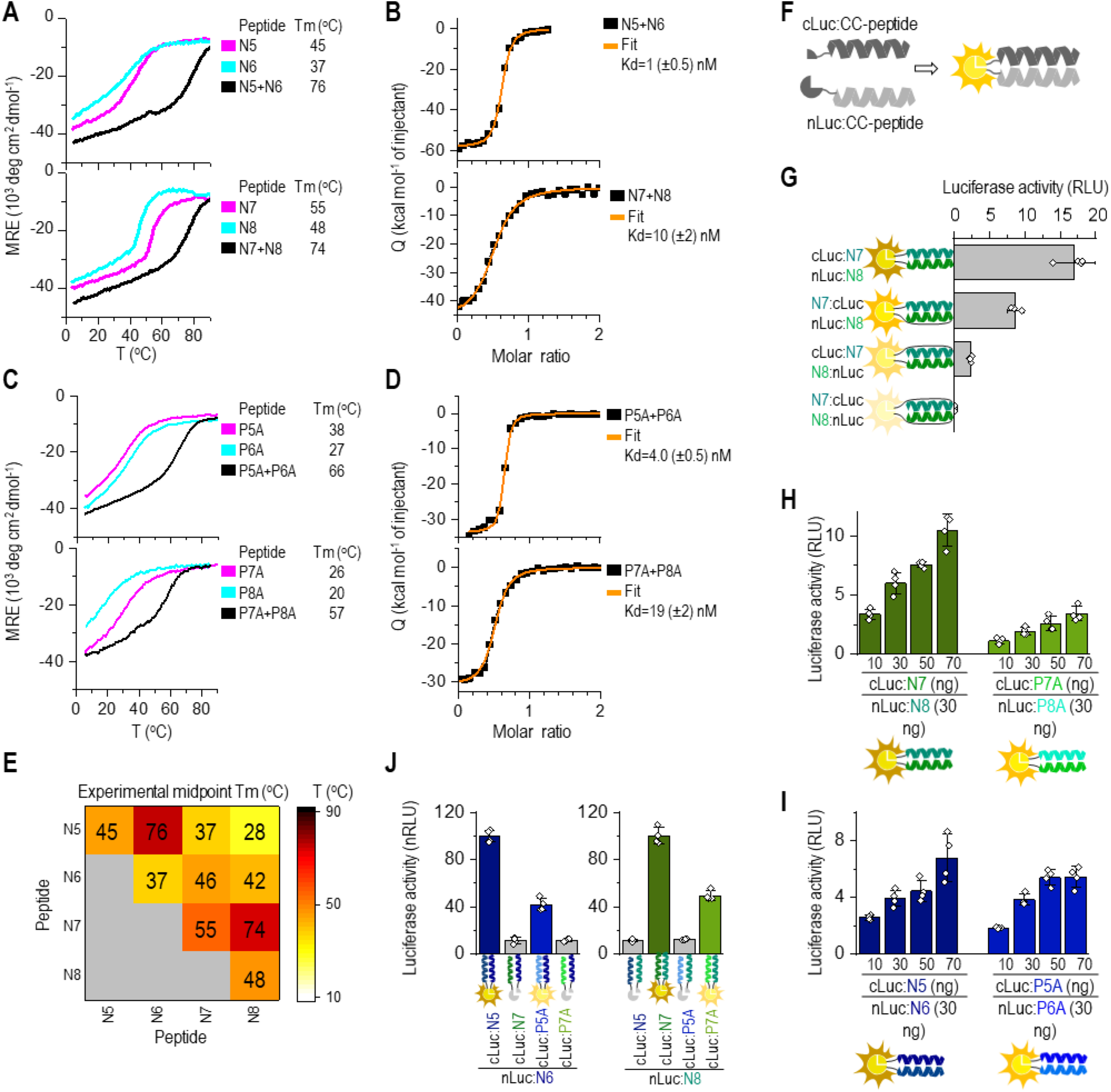
Orthogonal CC peptide pairs with K_d_ in the nanomolar range based on modifications at the *b*, *c*, and *f* sites. (**A,C**) Thermal denaturation profiles of peptides (40 μM; magenta and cyan) and CCs (20 μM each peptide; black) monitored by a CD signal at 222 nm. The midpoint Tm was calculated based on thermodynamic model fit ^33^. (**B,D**) Isothermal titration calorimetry (ITC) analysis of the binding affinity of designated CC peptide pairs. The binding isotherms of heat release per injection are depicted as a function of the increasing peptide-to-peptide molar ratio. The dissociation constant, K_d_^ITC^, was calculated using the two-state dimer association model. (**E**) Heat map of the matrix of the calculated midpoint Tm from thermal denaturation scans of all peptide combinations. (**F**) Scheme of reconstitution of CC-split luciferase managed by a CC-forming peptide pair. (**G**) Luciferase activity of reconstituted CC-split luciferase in HEK293T measured 48 h after transfection of HEK293T cells with a plasmid expressing a combination of nLuc tethered to N8 and cLuc tethered to N7 peptide. (**H,I**) Luciferase activity determined 48 h after transformation of HEK293T cells with plasmids expressing nLuc:N8 or nluc:N6; and cLuc tethered to N7 or P7A or cLuc tethered to N5 and P5A. (**J**) Orthogonality of designed N peptide set in HEK293T cells by co-cotransfection of the nLuc:N8 or nLuc:N6 fusion encoding plasmid, and cLuc:CC (CC stands for N5, P5A, N7, P7A). Reconstituted luciferase activity was measured 48 h after transfection. Amounts of plasmids are indicated in **Table S2**. The values (**G-J**) represent the means (± s.d.) from four independent cell cultures, individually transfected with the same mixture of plasmids, and are representative of two independent experiments.

In order to tune the stability of designed CCs and avoid homodimerization, we designed additional PA peptide pairs. The Ile residues at the *a* positions of the first or fourth heptad were replaced with Asn, resulting in peptide variants named P5A:P6A and P7A:P8A (PA peptides), respectively (**Table 1**; the structural model of P7A:P8A is depicted in **Fig S1A**). While the inclusion of an additional Asn residue destabilized the hydrophobic core ^28–30^, it also led to a decrease in the predicted helicity; thus, negatively contributing to the CC formation as confirmed experimentally. The CD spectra and thermal denaturation scans of the individual peptides showed that the second Asn within the PA peptide decreased their α-helical and thermal stability (**Fig S2C**). Furthermore, in contrast to the N variants, the individual PA peptides were present in solution in a monomeric state and formed dimers only when mixed with their cognate peptide, as shown by SEC-MALS analysis (**Fig S2D**). The Tm values of the P7A:P8A and P5A:P6A CC pairs were 64 °C and 57 °C, respectively (**Fig 1C)**. The ITC-measured Kd value was in the nanomolar range (**Fig 1D**). The orthogonality of PA peptides was confirmed by measuring the stability of all combinations of PA and N-type peptides (**Fig S2E**). The expected cognate peptide pairs exhibited higher Tm values than any other combination of peptides, indicating the preference of the peptides to bind to their designated partners.

### Heterodimeric CC pairs in mammalian cells for reconstitution of enzymatic activity

Characterization of CC pairs *in vitro* provided essential information about their stability. Next, the impact of a cellular environment on the formation of CCs between the cognate peptides was tested by the reconstitution of a split luciferase reporter in mammalian cells (scheme **Fig 1F**). The N- and C-terminal segments of split luciferase (henceforth, nLuc and cLuc, respectively) were genetically tethered to the N- and C-termini of the N8 and N7 peptides, respectively. All four combinations of CC–split luciferase reconstituted enzymatic activity in HEK293 cells (**Fig 1G, Fig S3A**). The highest activity was measured for the CC–split luciferases linked to the N-termini of cognate CC peptides; therefore, this setup was used for further experiments.

After establishing the most efficient setup of the CC-split luciferase, we analyzed the efficiency of CC formation for newly designed N-peptides (N5:N6 and N7:N8) and compared those with the PA-peptides (P5A:P6A and P7A:P8A; **Fig 1H**, **Fig 1I**). Both the N- and PA-peptide pairs reconstituted the CC-split luciferase; however, reconstitution of the PA-CC-split luciferases was less effective than the N-CC-split luciferase, which corroborated the *in vitro* stability results of the CC peptides’ stability, regardless of the homodimerization propensity of the N peptides. The orthogonality of all eight peptides was also tested in mammalian cells. In accordance with *in vitro* results, the nLuc:N6 paired only with cLuc:N5 or P5A but not with cLuc:N7 or P7A, and nLuc:N8 paired with cLuc:N7 and P7A (**Fig 1J**). The reconstitution of N-CC-split luciferase was more efficient compared to PA-CC-split luciferase, as reflected by the *in vitro* thermal stability of N- and PA-CCs, which makes N-type peptides very suitable for biological applications. While CD-based thermal denaturation and SEC-MALS cannot discriminate between the homo- and hetero-dimeric CCs, the reconstitution of CC–split proteins depends on the formation of CC heterodimers ^3^ and, therefore, confirms the heterodimerization of CC pairs.

### Tunable stability of CC pairs for biological applications

We have previously designed and reported an orthogonal, parallel NICP set of CC heterodimers ^17^ that contain the same electrostatic motifs as the N- and PA-peptides. Pairing CC-forming peptides with complementary electrostatic motifs (EKEK:KEKE and EKKE:KEEK at *e* and *g* positions) but different helical propensities at non-interacting *b, c,* and *f* heptad positions allows for facile fine-tuning of the stability of the resulting CC combinations ^16^. Consequently, pairing N-peptide variants with P variants (**Table S1** sequences) can create an extended range of pairs with different stabilities. Using the CC-split luciferase assay described above (**Fig 1F**), we analyzed the formation of N-peptides in combination with the P-peptides with a complementary electrostatic motif. NLuc was genetically tethered to the N-termini of the N8 or N6 peptide, and cLuc to N7, P7A, P7, P7SN, N5, P5A, P5, or P5SN peptide (**Fig 2A**, **Fig 2B**). The interaction matrix of all electrostatically complementary peptide combinations predicted the formation of CCs with a range of stability profiles, which were validated experimentally by thermal denaturation scans (**Fig S4A, Fig S4B**). The stability of CCs formed with the cognate N-and P-peptide pairs (Tm 43–68 °C) was lower compared to the N-type (Tm > 70 °C). We also investigated whether the stability of the cognate CC peptides with different helical propensities, determined *in vitro*, correlated with the reconstitution efficacy of CC-split luciferase in mammalian cells. The highest luciferase activity was determined for the combination of nLuc:N8 with cLuc:N7 and nLuc:N6 with cLuc:N5 (**Fig 2A**, **Fig 2B**; for CC-split luciferase see **Fig S3B**). In contrast, the CC-split luciferase combination, nLuc:N with cLuc:PSN pairs, generated the weakest signal. The results indicate that CC–split luciferase, as a reporter of CC stability in cells, adequately reflects the CC stability determined *in vitro*. As predicted, only orthogonal peptide variants with complementary electrostatic motifs reconstituted the functional luciferase (**Fig 2C**, **Fig 2D**). Furthermore, the wide range of CC stabilities obtained for all the pairs enables the use of different CC-dimerizing elements to fine tune the enzymatic activity.

**Figure 2.**
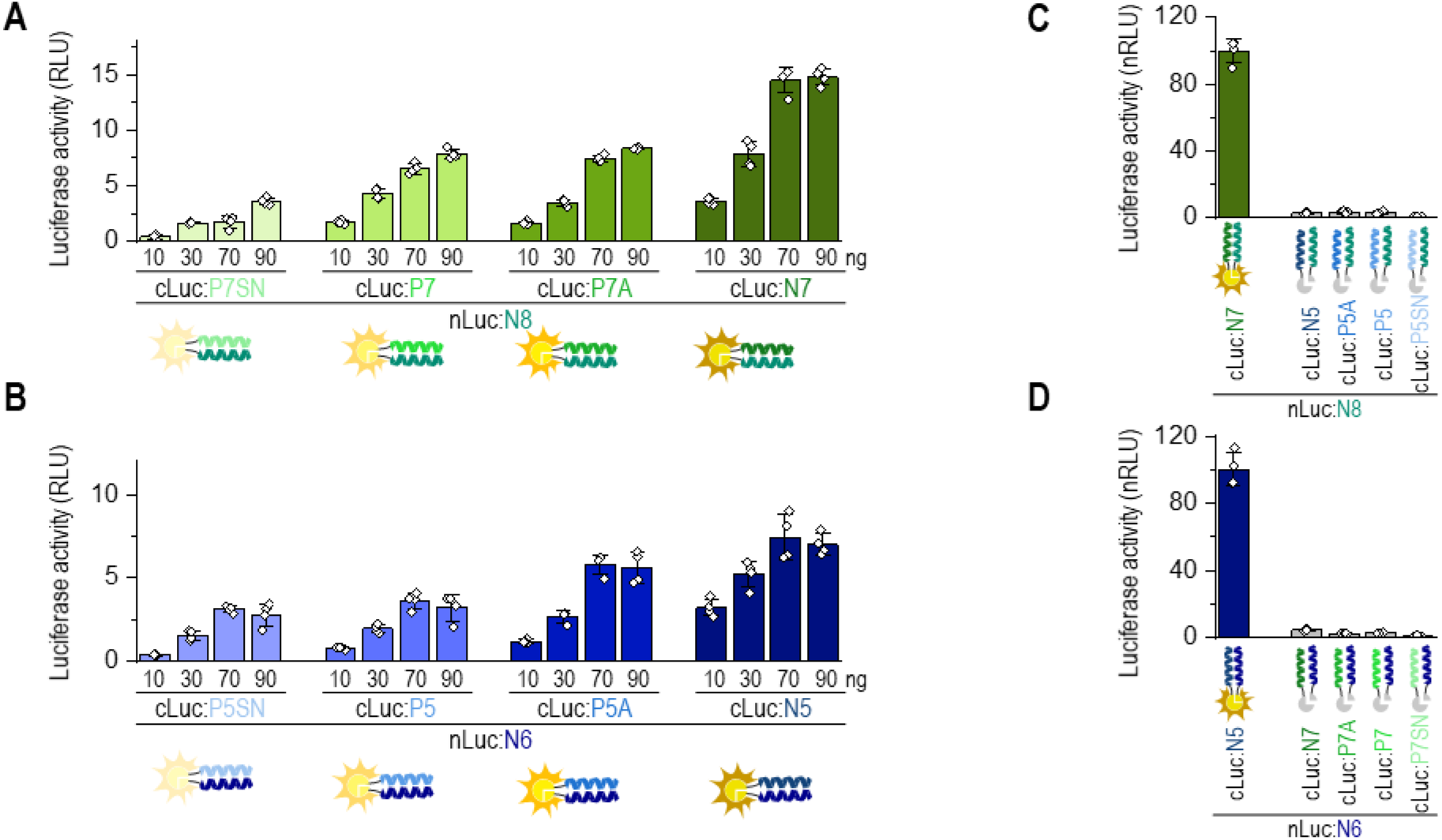
Reconstitution of split luciferase can be regulated by CC stability. Luciferase activity was determined 48 h after transformation of HEK293 cells with plasmids expressing nLuc:N8 or nluc:N6; and cLuc tethered to N7, P7A, P7, P7SN or to N5, P5A, P5, P5SN. (**A**, **B**) Increasing amounts of plasmids expressing cLuc:CC resulted in increased luciferase activity. **(C, D)** Only orthogonal peptide variants with complementary electrostatic motifs formed functional luciferase. The values represent the means (± s.d.) from four independent cell cultures, individually transfected with the same mixture of plasmids, and are representative of two independent experiments. Amounts of used plasmids are indicated in **Table S2**.

### Regulation of split luciferase activity in cells by a displacer peptide

The stability of CCs and their formation in mammalian cells can be tuned by combining CC-forming peptide pairs with different helical propensities. Thus, we examined whether the electrostatically cognate CC-forming peptide (CC–displacer peptide), with a higher helical propensity, could displace the weaker interacting peptide within CC in mammalian cells (scheme **Fig 3A**). CC–split luciferase was used as a reporter, and non-labeled peptides were used as competitors. Initially, we tested whether co-expressed CC–displacer peptide, N7, could compete with CC peptides tethered to the split luciferase (**Fig 3B**). Co-expression of the N7 CC–displacer peptide attenuated reconstitution of the split luciferase tethered to peptides N8:P7SN, N8:P7, or N8:P7A (displacer peptide N5 attenuated N6:P5 split luciferase activity; **Fig S5A**) but, as expected, failed to effectively compete with the immensely stable, N8:N7 CC (**Fig 3C**).

**Figure 3.**
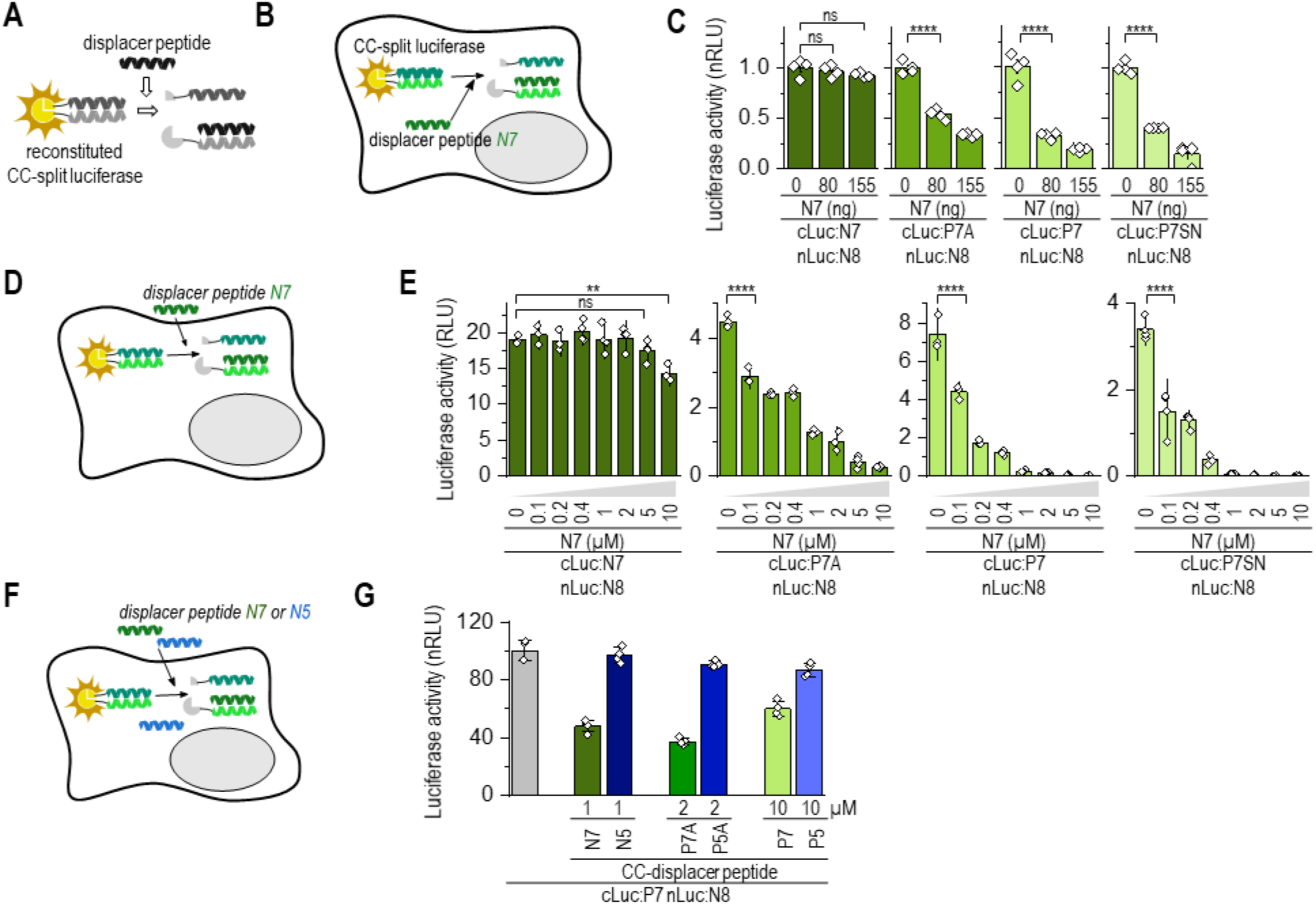
Modulation of split protein activity by competition based on CC stability. (**A**) Schematic representation of luciferase activity attenuated by a displacer peptide and (**B)** of attenuation of luciferase reconstitution in HEK293T cells with co-expressed peptide N7. Reconstitution of CC–split luciferase (nLuc:N8 and cLuc:CC) was attenuated by a cognate CC-displacer peptide (N7) that forms stronger CCs. (**C**) Luciferase activity of HEK293T cells co-transfected with plasmids expressing nLuc:N8 and cLuc tethered with N7, P7A, P7, or P7SN and a plasmid expressing N7 displacer peptide (0, 80, and 155 ng). Luciferase activity was measured 48 h after transfection. (**D**) Schematic representation of luciferase activity attenuated by a displacer peptide, which was added to split luciferase-expressing cells. (**E**) Luciferase activity of HEK293T cells expressing the CC-split luciferase treated with N7 peptide (0–10 μM in DOTAP) for 2 h. Forty-eight hours after transfection of HEK293T cells with plasmids expressing nLuc:N8 and cLuc tethered with N7, P7A, P7, or P7SN, cells were treated with CC-displacer peptide N7. **(F)** Schematic representation of luciferase activity attenuated by the addition of a displacer peptide with either matched or mismatched electrostatic motifs. **(G)** Luciferase activity of HEK293T cells treated with N7 or N5; P7A or P5A; P7 or P5 peptide (1, 2 or 10 μM in DOTAP) for 2 h. Cells were treated 48 h after transfection with plasmids expressing nLuc:N8 and cLuc:P7. The values (**C**, **E**, **G**) represent the means (± s.d.) from four independent cell cultures, individually transfected with the same mixture of plasmids, and are representative of two independent experiments. For the amounts of plasmids, see **Table S2**. Statistical analyses and the corresponding p-values are listed in **Table S3**.

Since the externally added peptide provides a more direct and potentially therapeutically relevant way to manipulate the activity of split proteins than co-expression of the CC–displacer peptide, we tested the efficacy of strand displacement by adding synthetic CC–displacer peptide to cells expressing reconstituted CC–split luciferase (**Fig 3D**). Three CC–displacer peptides with a cognate electrostatic motif (N7, P7A, and P7) were tested on four combinations of CC–split luciferase: N8:N7, N8:P7A, N8:P7, and N8:P7SN (**Fig 3E**, **Fig S5B**). As predicted, N7 displacer peptide, with its high-helical propensity, debilitated CC-split luciferase assembly most efficiently only 2 hrs after the addition of a synthetic displacer peptide (**Fig 3E**). The P7A, P7, and P7SN peptides, for which the midpoint Tm of N-P-CCs was more than 10 °C lower than N-CCs, failed to displace the cognate peptide from the N8:N7 split luciferase as efficiently (**Fig S5B**). Additionally, the N5 peptide effectively displaced cLuc:P5 from the N6:P5 split luciferase (**Fig S5C**).

Moreover, the CC-displacer peptide with mismatched electrostatic motif failed to displace the cognate CC peptide pair and, therefore, failed to attenuate the luciferase activity (**Fig 3G**, **Fig S5D**). Taken together, the results indicate that the formation and disruption of CC-mediated pairs can be regulated by competition with an electrostatically cognate peptide that forms stronger interactions.

### Transcriptional regulation by the strong CC dimers

In order to further test the utility of the designed coiled-coil set, we prepared CC-regulated transcriptional regulators. The transcription activator-like effector A (TALE-A) DNA-binding domain was fused to the N terminus of a cognate CC pair. VP16 activation domain ^35^ or Krüppel associated box (transcription repression domain; KRAB) ^36^, on the other hand, were fused to the C terminus of a CC-forming peptides (scheme **Fig 4A, Fig 4C**). The synthetic TALE-A DNA-binding domain ^37,38^ was linked to a single N8 and N6 peptide forming TALE-A:N8 and TALE-A:N6, respectively. The activation domain, VP16, was linked to N5, P5A, P5, N7, P7A, or P7 peptide, and the KRAB domain linked to N5, P5A, P5, or P5SN peptide. The transcriptional activator or repressor was reconstituted by combining TALE-A:CC with CC:VP16 and/or CC:KRAB. The reconstitution of the TALE-A DNA-binding domain and VP16 activation domain via CC heterodimeric interaction was determined by the expression of a reporter gene firefly luciferase.

**Figure 4.**
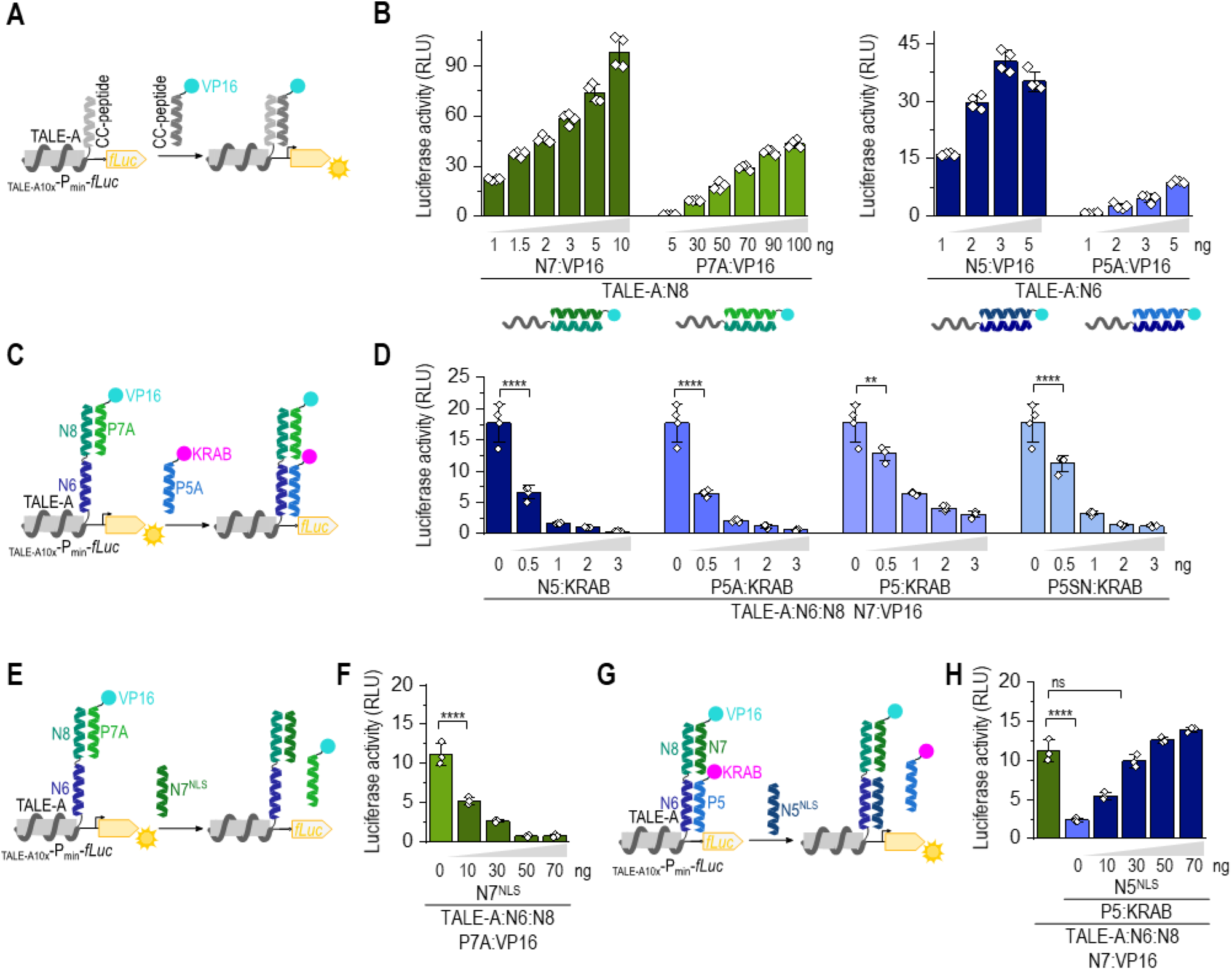
Tunable regulation of transcriptional activation by the selection of a CC partner peptide. (**A**) Schematic representation of CC–split transcription activator. Reconstitution via CC forms a functional transcription activator, which triggers the expression of a reporter located downstream of 10 TALE-A binding sites (_TALE-A10x_-P_min_-*fLuc*). (**B**) Reconstitution of CC–split transcription factor in HEK293T cells co-expressing TALE-A:N8; VP16 linked to N7 or P7A or TALE-A:N6 and VP16 linked to N5 or P5A; and the reporter luciferase (_TALE10x_-P_min_-fLuc). Luciferase activity was measured 48 h after transfection. (**C**) Schematic representation of CC-split transcription factor regulated by the addition of orthogonal CC-KRAB (suppressor) and CC-VP16 (activation) domain. (**D**) HEK293T cells were co-transfected with plasmids expressing TALE:N6:N8, VP16:N7, and KRAB linked to N5, P5A, P5, P5SN, and the reporter luciferase (_TALE10x_-P_min_-fLuc). Luciferase activity was measured 48 h after transfection. (**E**) Schematic representation of suppression of CC–split transcription factor (TALE:N6:N8-N7:VP16) activity by a CC–displacer peptide N7^NLS^. (**F**) Luciferase activity determined 48 h after transfection of HEK293T cells with plasmids expressing TALE:N6:N8, VP16:N7, CC–displacer peptide (N7^NLS^) and the reporter luciferase (_TALE10x_-P_min_-fLuc). (**G**) Schematic representation of CC–split transcription factor (TALE:N6:N8-N7:VP16-P5:KRAB) activity modulation with N5^NLS^ displacer peptide. (**H**) Luciferase activity determined 48 h after HEK293T cells were transfected with plasmids expressing TALE:N6:N8, VP16:P7A, P5:KRAB, N5^NLS^ and the reporter luciferase (_TALE10×_-P_min_-fLuc). The values (**B,D,F,G**) represent the means (± s.d.) from four independent cell cultures, individually transfected with the same mixture of plasmids, and are representative of two independent experiments. For amounts of plasmids, see **Table S2**. Statistical analyses and the corresponding p-values are listed in **Table S3**.

Initially, we tested a reconstitution of TALE-A:N8 or TALE-A:N6 with CC:VP16 (**Fig 4B**, **Fig S6A**). The highest expression of reporter luciferase under TALE promoter was detected for the combination of TALE-A:N8 with N7:VP16 and TALE-A:N6 with N5:VP16 (**Fig 4B**). As expected, VP16 tethered to a PA peptide variant with lower helical propensity resulted in a weaker transcription of the reporter. The efficiency of the reconstituted transcription factor was more dependent on the amount of the transcriptional activation-domain-(CC:VP16)-encoding plasmid than on DNA binding domain (TALE-A:CC), which can saturate the binding sites (**Fig 4B**, **Fig S6A**). Only the cognate CC peptide pairs (TALE-A:N6 with N5:VP16 and TALE-A:N8 with N7:VP16) triggered transcription of the reporter (**Fig S6B**). The results show that CC peptides with different stabilities can mediate the efficiency of transcriptional activation, with expression reflecting the stability of the CC pairs.

Orthogonal CC peptides provide a very effective means of constructing multifunctional proteins. In addition to genetically fused TALE-A to a single CC-peptide, we also created a construct, where the DNA binding domain was fused to two consecutive orthogonal CC-peptides (N6 and N8; scheme **Fig 4C**). In order to demonstrate the usability of orthogonal peptides, the TALE-A:N6:N8 was co-expressed with the CC:VP16 activator and/or CC:KRAB repressor domain. While the combination of TALE-A:CC with CC:VP16 acted as a transcriptional activator, the addition of CC:KRAB suppressed transcription (**Fig 4C**). Addition of the KRAB repressor tethered to CC N5, P5A, P5, or P5SN peptides (CC:KRAB), with different affinities to N6 of the reconstituted TALE-A:N6:N8 with N7:VP16, effectively suppressed transcription, correspondingly with their CC stability (**Fig 4D**). A relatively low amount of co-transfected CC:KRAB plasmid was required for the effective suppression of the transcription activator. Even when VP16 was replaced with the VPR activation domain, which is regarded as a stronger domain ^39^, the co-expressed CC:KRAB domain nevertheless suppressed the transcription (scheme **Fig S6D, Fig S6E**). The stability of CC formed between the CC:KRAB peptide and TALE-A:N6 determined the degree of transcription suppression, with P5SN:KRAB being the weakest suppressor. Therefore, the same DNA binding protein could be active, whether up- or down-regulated, which also represents a Boolean logic function B, NIMPLY A.

The activity of the multifunctional transcription factor, TALE-A:CC, could be regulated by CC:VP16 and CC:KRAB association as well as by the addition of a CC-displacer peptide. The same strategy of the CC-displacer peptide used for CC-split luciferase (**Fig 3**) could be used to regulate the repressor activity of TALE:N6:N8 with N7:VP16 (scheme **Fig 4E**) and P5:KRAB (scheme **Fig 4G**). The reconstitution of the CC–split transcription factor, similar to that of CC-split luciferase, was attenuated by co-expressed CC–displacer peptide tagged with a nuclear localization signal (NLS). The degree of attenuation reflected the stability of CC within the CC-transcription factor and CC-displacer peptide (**Fig 4F**, **Fig S6C**). As predicted, the N7 displacer peptide weakened the transcription activator assembly most efficiently. P7A and P7, with a weaker affinity toward N8, failed to prevent the reconstitution of a functional transcription as efficiently (**Fig S6D**). Besides, through the addition of increasing amounts of N5^NLS^ peptide (cognate to N6), we were able to displace the KRAB suppressor domain and regain transcriptional activity (**Fig 4G, 4H**). Altogether, we demonstrated that the array of orthogonal CC peptide pairs that form CCs with various ranges of stability represents an effective tool for rapid and effective regulation of biological activity, likely transferrable to other enzymes, assemblies, and processes.

### CC-mediated reporters for viral fusion protein-mediated syncytium formation

We realized that the high affinity between CC domains could be used to monitor the fusion of cells for the syncytium formation. SARS-CoV-2 entry into host cells is mediated by a viral spike (S) protein, which binds to the human Angiotensin-converting enzyme 2 (ACE2) as a viral entry receptor ^40^. Viral entry depends on the binding of the S1 domain of spike protein to the ACE2 receptor, which facilitates viral attachment to the surface of target cells. Virus entry also requires the processing of the spike protein by the host transmembrane serine protease 2 (TMPRSS2), which exposes the S fusion peptide. This process enables the fusion of the viral and cellular membranes, a process driven by the spike protein’s S2 subunit ^24,41^. Monitoring membrane fusion triggered by the binding of a viral fusion protein to the receptor represents an important assay for monitoring this key process in viral infections. Many inhibitors, as well as vaccines, target viral attachment to host cells and the fusion process. Experiments using active viruses or pseudoviruses are labor-intensive, slow, and require specific safety measures. Fluorescent proteins and transcriptional activation in target cells have been used to monitor fusion triggered by the S protein of SARS-CoV-2 with ACE2 receptor ^40^. An assay was prepared based on the fusion of cells expressing ACE2 receptor protein (acceptor cells) and cells expressing the S protein (donor cells) at their surfaces. Formation of cell fusion was detected by co-transfection of two cell groups with split luciferase reporters (cLuc:P7A and nLuc:P8A or cLuc:N7 and nLuc:N8) with ACE2 and SARS-CoV-2 S protein, respectively (**Fig 5A**). Mixing of donor and acceptor cells resulted in a rapid reconstitution of the reporter in fused cells. The addition of CCs to a split luciferase reporter acts as a strong dimerization domain, generating a high signal following cell fusion, but only for cell combinations that expressed S protein and hACE2 (**Fig 5B**).

**Figure 5.**
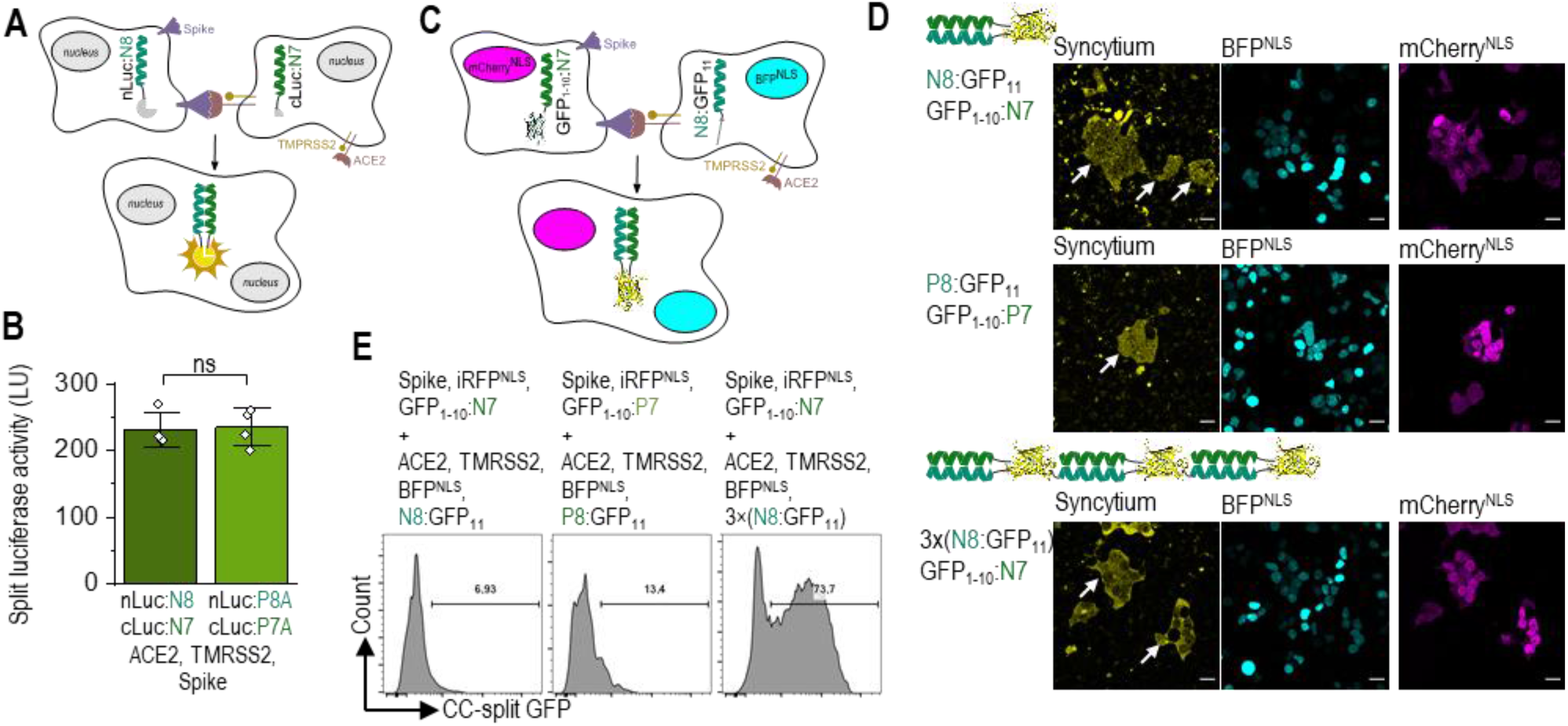
CC-split enzymes as reporters for viral fusion protein-mediated syncytium formation. (**A**) Schematic representation of S protein-mediated cell-cell fusion assay. The donor cell is identified as the cell co-expressing CoV-2 Spike protein and nLuc:N8, the acceptor cells with co-expression of ACE2 receptor, TMPRSS2, and cLuc:N7. Syncytium formation is detected with reconstituted CC-split luciferase activity. (**B**) Luciferase activity as an indicator of cell fusion. 24 h after transfection, donor HEK293T cells transfected with plasmids expressing nLuc:N8 or nLuc:P8A, CoV-2 Spike-protein, and acceptor cells expressing cLuc:N7 or cLuc:P7A, ACE2, TMPRSS2 were mixed in 1:1 ratio. Luciferase activity was measured 3 h later. The values represent the means (± s.d.) from four independent cell cultures, individually transfected with the same mixture of plasmids, and are representative of two independent experiments. (**C**) Schematic representation of syncytium formation between the donor cells expressing CoV-2 Spike protein, CC-split GFP and mCherry^NLS^, and the acceptor cells expressing ACE2 receptor, TMPRSS2, CC-split GFP, and BFP^NLS^. The fused cells are identified with BFP and mCherry nuclei and reconstituted CC-split GFP. (**D**) Confocal microscopy images of a mixture of donor HEK293T cells expressing the CoV-2 Spike protein, mCherry^NLS^, and GFP_1-10_:N7 or GFP_1-10_:P7 and the acceptor cells expressing ACE2 receptor, TMPRSS2, BFP^NLS^, and N8:GFP_11_, P8:GFP_11_ or 3×(N8:GFP_11_). After 3 h mixing of donor and acceptors cells, syncytia (indicated with an arrow) were observed by a reconstituted GFP (yellow) and co-localization of BFP (cian) and mCherry (magenta). (**E**) Flow cytometry analysis of a mixture of donor cells expressing the CoV-2 Spike protein, iRFP^NLS^, and GFP_1-10_:N7 or GFP_1-10_:P7 and the acceptor cells expressing ACE2 receptor, TMPRSS2, BFP^NLS^, and N8:GFP_11_, P8:GFP_11_ or 3×(N8:GFP_11_). Percent of reconstituted split GFP for double iRFP and BFP positive cells 3 h after mixing donor and acceptor cells. For gating strategy, see **Fig S7B**. Representative results of two independent experiments are shown. For amounts of plasmids, see **Table S2**. Statistical analyses and the corresponding p-values are listed in **Table S3**.

In addition to a CC-split luciferase reporter, we set out to design a CC-split green fluorescent protein (GFP) reporter ^42^, which enables syncytium visualization via confocal microscopy and quantification using flow cytometry. The fusion of CC peptides to the split GFP improves the efficiency of the reporter reconstitution and enables the detection of syncytia. We designed GFP_1-10_ genetically fused to the N7 or P7 peptide (GFP_1-10_:N7, GFP_1-10_:P7) and GFP11 fused to N8 or P8 peptide (N8:GFP_11_, P8:GFP_11_; **Fig 5C**). The donor cells expressed S protein, iRFP or mCherry, and CC-split GFP (GFP_1-10_:N7 or GFP_1-10_:P7). The acceptor cells expressed ACE2 receptor, BFP, and CC-split protein. Using CC-split GFP, we were able to detect cell-cell fusion both utilizing a confocal microscope and via flow cytometry (**Fig 5D, 5E)**. All syncytia also expressed iRFP and BFP. In order to increase the sensitivity of flow cytometry for syncytium detection, a construct with linked three repeats of N8:GFP11 (3×(N8:GFP_11_)) was designed (**Fig 5D**), which enables three molecules of GFP to reconstitute in a fluorescent reporter. Indeed, this kind of reporter proved to be even more effective compared to the construct containing only a single CC-forming peptide (**Fig 5E**).

The S protein-/ACE-mediated fusion process could, thus, be monitored at different stages, and inhibitor efficiencies of this process could be detected. We demonstrated the effect of inhibiting TMPRSS2 protease that is required to cleave S protein and expose the fusion protein of spike protein prior to fusion. Camostat mesylate ^43^ has been previously tested as an inhibitor in the active viral assay^43^ (**Fig 6A**). Cell fusion was detectable without the presence of TMPRSS2 protease (**Fig S7A**); however, a strong signal increase was observed upon the co-expression of the protease (**Fig. 6B),** which confirms the role of TMPRSS2 in CoV-2 spike-protein-mediated syncytia formation as previously reported ^25^. Additionally, fusion could be inhibited by blocking the ACE2 receptor by the addition of a soluble RBD domain ^44,45^ (**Fig.6C, 6D**). Increasing amounts of RBD protein indeed reduced the number of syncytium cells generated by restraining ACE2 and CoV-2 S protein interaction **(Fig 6D**). Inhibition of cell-cell fusion was also detected using flow cytometry.

**Figure 6.**
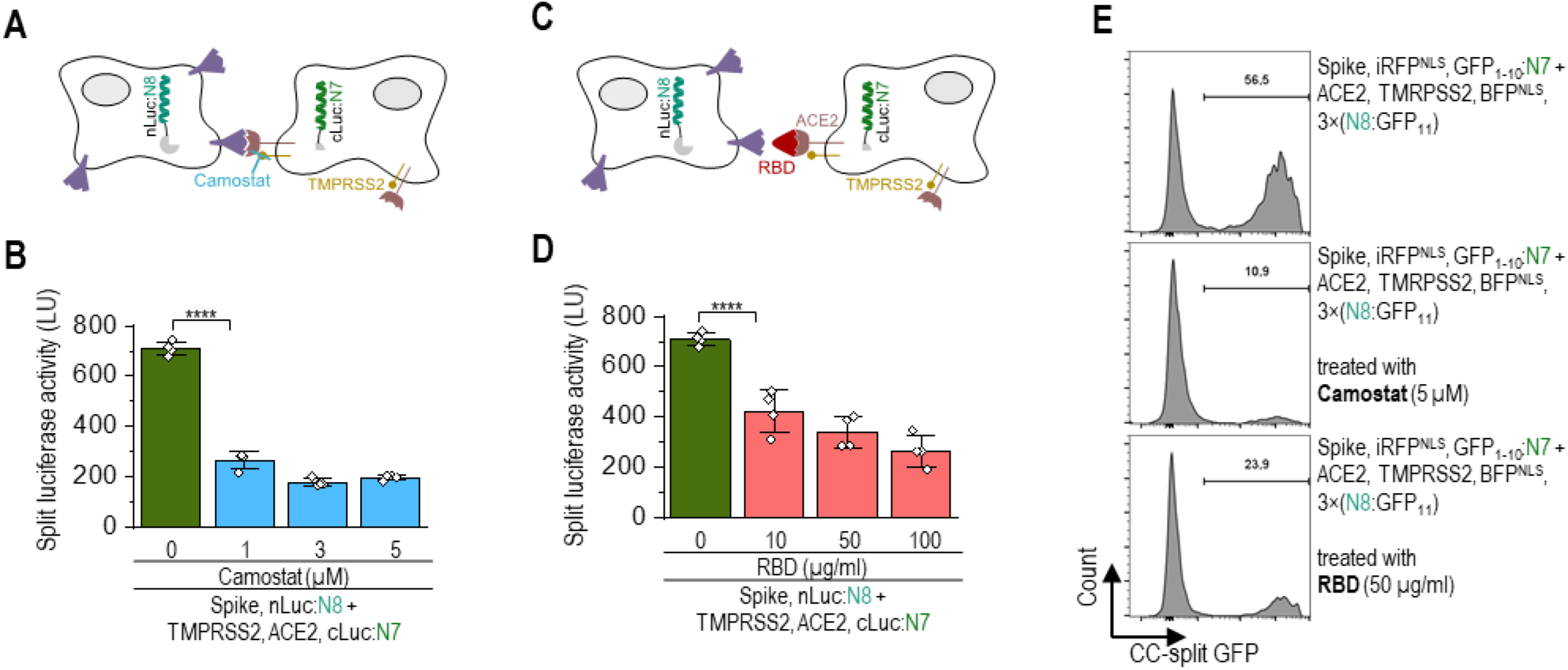
Inhibition of Spike-protein hACE2-mediated fusion determined by the CC-reporter assay. (**A,C**) Schematic representation of inhibition of SARS CoV-2 S protein-mediated cell-cell fusion by TMPRSS2 protease inhibitor Camostat (**A**) and soluble RBD protein domain binding to the ACE2 receptor (**B**). (**B,D**) Luciferase activity of a mixture of donor and receptor cells treated with Camostat (**B**) or RBD (**D**). Donor cells were transfected with plasmids expressing nLuc:P8A, CoV-2 S protein, and the acceptor cells expressed cLuc:P7A, ACE2, TMPRSS2. The donor and acceptor cells were mixed and treated with RBD or Camostat. Luciferase activity was measured 3 h later. The values (B,D) represent the means (± s.d.) from four independent cell cultures, individually transfected with the same mixture of plasmids, and are representative of two independent experiments. (**E**) Flow cytometry analysis of a mixture of donor cells expressing the SARS CoV-2 S protein, iRFP^NLS^, and GFP_1-10_:N7 or GFP_1-10_:P7 and the acceptor cells expressing ACE2 receptor, TMPRSS2, BFP^NLS^, and N8:GFP_11_, P8:GFP_11_ or 3×(N8:GFP_11_). Percent of reconstituted split GFP for double iRFP and BFP positive population 3 h after mixing donor and acceptor cells treated with Camostat or RBD. For gating strategy, see **Fig S7B**. Representative results of two independent experiments are shown. For amounts of plasmids, see **Table S2**. Statistical analyses and the corresponding p-values are listed in **Table S3**.

## Discussion/Conclusion

In this study, we designed and explored the potential of CC–peptide pairs with high stability as a tool for biological applications. We expanded the validated orthogonal set of CC dimer-forming peptides ^9^ with two *de novo* designed orthogonal CC–peptide pairs that exhibit low nanomolar Kd values and a Tm around 70 °C. This improved stability was achieved by increasing the helical propensity at noncontact positions and by addition of a single Asn residue at *a* position. Since the modifications did not alter the dimerization interface, pairing specificity remained unaffected ^9^. A single Asn residue (per peptide) that supports the desired orientation was positioned at the first or fourth heptad. Although a previous study by Fletcher et al. ^28^ showed that a single Asn residue positioned at the end of the heptad had a minor impact on stability, we determined that the peptide pairs, P7A:P8A and P5A:P6A, with two Asn residues per peptide, had lower stability but still in the nanomolar range, and the Tm values were reduced by at least 10 °C. Contrastingly, the introduction of an additional Asn residue at the *a* position of N-type CC peptides prevented homodimerization in the P7A:P8A pair and reduced the helical content of individual peptides.

The application of newly designed CCs in a cellular environment confirmed their high affinity in the cellular milieu and enabled regulation of activity of enzymes and transcriptional activation through genetic fusion with the CC-forming peptides. The described peptide set enabled fine-tuning of the affinity by varying the helical propensity of cognate peptide pairs while maintaining the electrostatic pattern. In comparison to the previously used set in mammalian cells^2^, new pairs are more stable and were used to demonstrate new modalities of regulation through the displacement of the weaker peptide from the existing CC pair, enabling up- or down-regulation with a wide, dynamic range with the same promoter. Genetic fusion of two orthogonal CCs to the DNA binding domain enabled transcription repressor and activator design within a single molecule. Transcriptional activity of the TALE-A:CC DNA-binding domain and CC:VP16 was suppressed by the addition of a CC:KRAB domain. Also, externally added peptides could regulate activity within cells.

Finally, high-affinity CC pairs were used to set up the assay for monitoring cell fusion. Genetic fusion to protein interaction domains can guide protein complex formation ^46^, and strong affinities are needed for a robust output in comparison to relying solely on the reconstitution of split GFP ^43^. We established an assay to monitor the SARS-CoV-2 spike-protein-mediated cell-cell fusion, which allowed for quantitation by reconstitution of a CC-split luciferase reporter or visualization of syncytia by the reconstituted CC-split GFP. The split luciferase reporter genetically fused to CCs produced a robust response upon syncytia formation, guided by the interaction between SARS-CoV-2 spike protein and ACE2 receptor. Addition of the TMPRSS2 inhibitor, Camostat, or RBD, which targets different steps in the fusion process, inhibited syncytia formation, validating the proteinase in S protein processing and binding to ACE2 receptor as targets for viral entry. The strategy of self-reconstituted split GFP previously described by Kamiyama et al. ^42^ was used to further intensify the generated fluorescent signal so it could detect formed syncytia upon mixing donor and acceptor cells, enabling better visualization and flow cytometric analysis. This rapid (~ 3hrs), sensitive assay enables screening of the inhibitors of this process as potential drugs to prevent viral infection. In conclusion, high-affinity CC dimers represent valuable tools for inquiring and regulating many biological processes.

## Methods

### Peptides, plasmids, and cell lines

The peptides used in this study (Proteogenix, France) were protected at the N- and C-termini by acetylation and amidation, respectively. All peptides were dissolved in a stock concentration of approximately 1 mg/ml in deionized water, except for the water-insoluble N1 peptide, which was dissolved in 10 % (w/v) ammonium bicarbonate. The exact peptide concentrations were determined based on absorbance at 280 nm, and the extinction coefficients were calculated by the ProtParam web tool (https://web.expasy.org/cgi-bin/protparam/protparam). All plasmids (listed in **Table S4**) were constructed using the Gibson assembly method. A human embryonic kidney cell line, HEK293T (ATCC CRL-3216), was cultured in complete media (DMEM; 1 g/l glucose, 2 mM L-glutamine, 10 % heat-inactivated FBS (Gibco)) with 5 % CO_2_ at 37 °C. We used plasmid pcDNA3 (Invitrogen) to express CCs and its fusions with split enzymes. *Renilla* luciferase (phRL-TK, Promega) was used as a transfection control in all experiments in mammalian cells.

### Immunoblotting

HEK293T cells were seeded in 6-well plates (Techno Plastic Products) at 2.5 × 10^4^ cells per well (1 ml). The next day, the cells were transiently transfected with plasmids expressing CCs or CC-split luciferases. The total amount of DNA for each transfection was 2.5 μg. Then, 48 h after transfection, the cells were washed with 1 ml PBS and lysed in 100 μL of lysis buffer (40 mM Tris-HCl, pH 8.0, 4 mM EDTA, 2 % Triton X-100, 274 mM NaCl with a cocktail of protease inhibitors (Roche)). Cells were incubated on ice for 10 min and then centrifuged for 15 min at 17,400 rpm at 4 °C to remove cell debris. The total protein concentration in the supernatant was determined using a BCA assay. Proteins from the supernatant were separated on 15 % SDS–PAGE gels (120 V, 60 min) and transferred to a nitrocellulose membrane (350 mA, 30 min). Membrane blocking, antibody binding, and membrane washing were performed with an iBind Flex Western device (Thermo Fisher) according to the manufacturer’s protocol. The primary antibodies were mouse anti-Myc (Cell signaling technologies; diluted 1:2,000), rabbit anti-HA (Sigma H6908; diluted 1:2,000), and mouse anti-β-actin (Cell Signaling 3700; diluted 1:2,000). The secondary antibodies were HRP-conjugated goat anti-rabbit IgG (Abcam ab6721; diluted 1:3,000) and HRP-conjugated goat anti-mouse IgG (Santa Cruz, sc2005; diluted 1:3,000). The secondary antibodies were detected with Pico or Femto Western blotting detection reagent (Super Signal West Femto; Thermo Fisher) according to the manufacturer’s protocol.

### CD spectroscopy

CD measurements were performed with a ChiraScan instrument (Applied Photophysics, UK) equipped with a Peltier thermal control block (Melcor, NJ, Laird Technologies). The final concentrations of individual peptides were 40 μM in Tris buffer (50 mM Tris-HCl, pH 7.5, 150 mM NaCl, and 1 mM TCEP). The mixture also contained a 20 μM concentration of each peptide. All experiments were performed in a quartz cuvette (Hellma, Germany) with a path length of 1 mm. CD spectra were measured every 1 nm from 200–280 nm with a bandwidth of 1 nm and 0.5 s integration time. The results are the average of three scans. Temperature denaturation scans were performed by measuring the CD signal at 222 nm from 0 to 95 °C in 1 °C increments at a rate of 1 °C/min. The actual sample temperatures were measured directly in the cuvette using a temperature probe. The CD spectrum was measured at 95 °C and at 20 °C after rapid cooling. For all peptides, the spectra after refolding matched the ones taken before the temperature scan.

### SEC-MALS

A Waters e2695 HPLC system coupled with a 2489 UV detector (Waters, MA) and a Dawn 8+ multiple-angle light scattering detector (Wyatt, CA) was used for SEC-MALS experiments. Peptides (200 μM in 50 mM Tris-HCl with a pH of 7.5, 150 mM NaCl, and 10 % glycerol) were filtered using Durapore 0.1 μm centrifuge filters (Merck Millipore, Ireland) before being injected onto the Biosep SEC-S2000 column (Phenomenex, CA), with the exception of the peptide N2, which was injected in a Superdex 30 increase column (GE, IL). The mobile phases used for the separations were 50 mM Tris-HCl, pH 7.5, 150 mM NaCl and 50 mM Tris-HCl, pH 7.5, 500 mM NaCl for the Biosep SEC-S2000 column and the Superdex 30 increase column, respectively. The injection volume was 50 μl, and the flow rate was 0.5 ml/min. Data analysis was carried out with Astra 7.0 software (Wyatt, CA) utilizing the theoretical extinction coefficient calculated by ProtParam.

### ITC

MicroCal VP-ITC (Microcal, Northampton, MA) was used to measure the equilibrium dissociation constant (K_D_) of orthogonal CC pairs. A 280-μL syringe was filled with the peptide solution (10 μM in 50 mM Tris-HCl with a pH of 7.5, 150 mM NaCl, and 1 mM TCEP). The samples were injected into an ITC cell (volume 1.3862 ml) filled with a complimentary peptide (1 μM) in a matching buffer through a 27-step process. The injection volumes were 10 μl and were spaced 600 s apart, with the exception of the first priming injection of 2 μl with 3200 s spacing. Before each ITC experiment, the sample and buffer solutions were degassed in an ultrasonic water bath. The integration of heat pulses and model-fitting were performed with Origin 7.0 using a single-site binding model.

### Modeling CC interactions and structures

To evaluate the orthogonality of the designed CC sets, the interactions between all possible peptide pairs were predicted by the scoring function introduced by Potapov et al. ^31^. For each peptide pair, the interaction score was calculated for three different relative alignments of the peptide chain. The amino acid sequences were either completely aligned or staggered by one heptad in the reverse or forward direction. The lowest score was taken as the interaction score of the peptide pair.

The Potapov scoring function ignores the contribution of amino acids at the *b*, *c*, and *f* positions and, therefore, cannot be used to predict the impact of mutations in these positions on CC stability. The scoring function introduced by Drobnak et al. ^16^ was used to evaluate the effect of *b*, *c*, and *f* mutations on CC stability. The melting temperatures for all peptide pairs were calculated using the following equation:

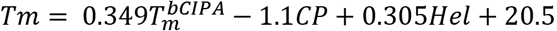

where 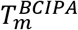 is the melting temperature predicted by bCIPA ^47^; CP is the peptide pair charge product; and *Hel* is the number of residues in a peptide pair with helicity above 0.05, as predicted by Agadir ^32^.

The model structures of designed CCs were built using the ISAMBARD modeling package ^48^. Structural parameters (superhelix radius, pitch, and Crick angle) were optimized using a genetic algorithm. The following space of parameters was searched: superhelix radius 5 ± 1 Å, pitch 200 ± 60 Å, Crick angle at *a* position 26 ± 27°. The relative shift along the superhelical axis was assumed to be zero. The internal energy was evaluated using the BUDE force field ^49^.

### Split luciferase and split transcription factor reconstitution

HEK293T cells were seeded in 96-well plates (Corning) at 2.5 × 10^4^ cells per well (0.1 ml). The next day, cells were transiently transfected with plasmids expressing CCs fused to split luciferase or plasmids expressing CC-split transcription factor (sequences are shown in **Table S4**), and phRL-TK (Promega) constitutively expressing *Renilla* luciferase (5 ng per well, for normalization of transfection efficiency) using the PEI transfection reagent. The total amount of DNA for each transfection was kept constant by adding appropriate amounts of the control plasmid pcDNA3 (Invitrogen).

For displacement of CCs by peptides, the HEK293T cells were transfected with plasmids expressing N- and C-terminal segments of split luciferase (nLuc and cLuc) fused to CC-forming peptides or CC-split transcription factors, plasmids expressing CC-displacer peptides under cytomegalovirus (CMV), and phRL-TK (Promega) constitutively expressing *Renilla* luciferase (5 ng per well) using the PEI transfection reagent Synthetic peptides were transfected into cells using DOTAP transfection reagent according to manufacturer instructions. Briefly, 48h after plasmid transfection, peptides were diluted in Hepes buffer to the working concentration of 100μM and then further diluted to final concentrations used on cells. DOTAP transfection reagent was added to peptide solution and, after 15-minute incubation, transferred to cell media. Two hours later, the media was removed, cells were washed with PBS to remove any additional peptides in media and lysed using 1x Passive lysis buffer.

### Viral fusion protein-mediated syncytium formation –Luciferase assay

Syncytia is formed by the fusion of cells expressing SARS-CoV-2 S protein on the cell surface with neighboring cells expressing ACE2 receptor leading to the formation of multi-nucleate enlarged cells. For fusion assay, the spike expressing (donor) cells and ACE2 receptor-expressing (acceptor) cells were mixed in 1:1 ratio. HEK293T cells were seeded in 6-well plates (Corning) at 5 × 10^4^ cells per well (2 ml). The next day, cells were transiently transfected with plasmids. The acceptor group was transfected with plasmids encoding cLuc:CC, ACE-2 receptor, and TMPRSS-2. The donor group was transfected with nLuc:CC and SARS-CoV-2 S protein using the PEI transfection reagent. The total amount of DNA for each transfection was kept constant by adding appropriate amounts of the control plasmid pcDNA3 (Invitrogen). 24h after transfection, cells were detached from 6 well plates using a solution of PBS, EDTA (2,5mM). Wells were additionally washed with DMEM, 10% FBS. Cells were then centrifugated (5min, 1500rpm), resuspended in 1ml DMEM, 10%FBS, and counted using the automated cell counter, Countess. Both groups of resuspended cells were diluted to a final concentration of 1×10^5^ cells/ml, mixed in ratio 1:1, and seeded to 96-well plates. TMPRSS2 inhibitor Camostat mesylate (1-5μM) or RBD (10-100μg/ml) were added to diluted cells expressing ACE2 receptor before mixing them with cells expressing S protein.

Three hours after mixing both cell groups, luciferase activity was measured as previously described ^3^, 10 μl of 0.5mM luciferase substrate luciferin and 2.5mM ATP was added per well before luciferase activity was measured with Orion II microplate reader, Berthold Technologies.

### Viral fusion protein-mediated syncytium formation-Flow cytometry

For flow cytometer analysis, the cells were seeded in a 12-well plate (Costar) (1 ml, 2.5 × 10^5^ cells/ml). At 70% confluency, the donor HEK293T cells were transfected with plasmids expressing S protein, fluorescent protein, iRFP^NLS^, and syncytia reporter split fluorescent protein, GFP1-10 (GFP_1-10_:N7, GFP_1-10_:P7 or GFP_1-10_). Acceptor cells were transfected with plasmids expressing a fluorescent protein, BFP^NLS^, ACE2 receptor, and syncytia reporter split protein, GFP_11_ (N8:GFP_11_, 3×(N8:GFP_11_), P8:GFP_11_)using PEI transfection reagent. Plasmid pcDNA3 was used to adjust the amount of plasmid DNA. Eighteen hours post-transfection, the spike expressing cells were detached from the surface using PBS, EDTA (2.5mM). Wells were additionally washed with DMEM, 10% FBS. Donor cells were then added to the ACE2 acceptor cells in 1:1 ratio. Syncytia were analyzed 3 h later. For inhibition of syncytium formation, soluble RBD (50-100μg/ml) or Camostat (50-100μM) was added to the cells at the same time as a mixture of cells was formed.

Syncytia formation was analyzed using the spectral flow cytometer Aurora with the blue, violet, and red lasers (Cytek). The cells gated positive for iRFP and BFP were examined for fluorescence intensity of reconstituted split GFP. The fluorescence intensity is presented as the median value. Data were analyzed using FlowJo (TreeStar, Ashland, OR, USA) and SpectroFlo (Cytek) software.

### Viral fusion protein-mediated syncytium formation-confocal microscopy

For confocal microscopy, acceptor cells were seeded in an 8-well tissue culture chamber (μ-Slide 8 well, Ibidi Integrated BioDiagnostics, Martinsried München, Germany) (0.2 ml, 2.5 × 10^5^ cells/ml), and acceptor cells were seeded in 24-well plates. At 70% confluency, the donor HEK293T cells were transfected with plasmids expressing S protein, syncytia reporter split fluorescent protein GFP1-10 (GFP_1-10_:N7, GFP_1-10_:P7 or GFP_1-10_) and fluorescent protein mCherry^NLS^, and the acceptor cells were transfected with plasmids expressing the ACE2 receptor, syncytia reporter split protein GFP11 (N8:GFP_11_, 3×(N8:GFP_11_), P8:GFP_11_), and fluorescent protein BFP using PEI transfection reagent. Plasmid pcDNA3 was used to adjust the amount of plasmid DNA. Eighteen hours post-transfection, the spike expressing cells were detached from the surface using EDTA (2,5mM) and added to the ACE2 acceptor cells in 1:1 ratio. Syncytia were analyzed 3 h later.

Microscopic images were acquired using the Leica TCS SP5 inverted laser-scanning microscope on a Leica DMI 6000 CS module equipped with an HCX Plane-Apochromat lambda blue 63 × oil-immersion objectives with NA 1.4 (Leica Microsystems, Wetzlar, Germany). A 488-nm laser line of a 100-mW argon laser with 10% laser power was used for split GFP excitation, and the emitted light was detected between 500 and 540 nm. A 1-mW 543-nm HeNe laser was used for mCherry:N6:NLS transfection control excitation and emitted light was detected between 580 and 620 nm. A 50-mW 405-nm diode laser was used for BFP transfection control, and emitted light was detected between 420 and 460 nm. The images were processed with LAS AF software (Leica Microsystems) and ImageJ software (National Institute of Mental Health, Bethesda, Maryland, USA).

### Dual-luciferase assays

At indicated time points, the cells were lysed in Passive Lysis 1× Buffer (Promega) and analyzed with a dual-luciferase reporter assay to determine the firefly luciferase and the *Renilla* luciferase activities (Orion II microplate reader, Berthold Technologies). Relative luciferase units (RLU) were calculated by normalizing the firefly luciferase value to the constitutive *Renilla* luciferase value in each sample. Normalized RLU values (nRLU) were calculated by normalizing the RLU values of each sample to the value of the indicated sample within the same experiment.

### Software and statistics

Graphs were prepared with Origin 8.1 (http://www.originlab.com/), and GraphPad Prism 5 (http://www.graphpad.com/) was used for statistical purposes. An unpaired two-tailed t-test (equal variance was assessed with the F-test, assuming normal data distribution) was used for statistical comparison of the data. If not otherwise indicated, each experiment was independently repeated at least three times. Each experiment was performed in at least three biological parallels.

## Data availability

The authors declare that the data supporting the findings of this study are available within the paper and its supplementary information files. The raw data are available from the corresponding author upon reasonable request.

## Abbreviations

CC: coiled coil
CCPO: coiled-coil protein origami
fLuc: firefly luciferase
rLuc: renilla luciferase
nLuc: N-terminal half of firefly luciferase
cLuc: C-terminal half of firefly luciferase
TALE: DNA-binding domain of TALEN
TALE10x: a TALE binding element concatenated ten times
Kd: dissociation constant
RLU: relative light units
Tm: denaturation temperature
s.d.: standard deviation
CD: circular dichroism
MRE: mean residual ellipticity
ITC: isothermal titration calorimetry
SEC: size-exclusion chromatography
SEC-MALS: SEC-multi angle light scattering

## Author contributions

T.P. cloned, purified, and experimentally characterized the coiled coils. P.D. helped with cloning and transfection experiments. J.A. designed the peptides. F.L. and T.P. performed the SEC-MALS experiments. M.B. and T.P. analyzed the experimental data. M.B. and R.J. led the research, designed the experiments, and wrote the manuscript. All authors discussed the results and reviewed and contributed to the manuscript.

## Acknowledgements

We thank Stefan-u Pöhlmann for pGC vectors encoding human ACE2 receptor and virus SARS −2 spike protein. We thank Sanjin Lulić for cloning, Ernest Špranger for protein isolation, dr. San Hadži for help with the CD measurements and Hana Esih and Tina Fink for RBD production and isolation. The research was supported by the Slovenian Research Agency (research core funding no. P4-0176 and J3-7034, J1-9173).

## Supporting Information

