## Supplemental material for "Coiled-coil heterodimers with increased stability for cellular regulation and sensing SARS-CoV-2 spike protein-mediated cell fusion"

Short: **Highly stable heterodimeric parallel coiled coils**

Tjaša Plaper[1,3], Jana Aupič[1], Petra Dekleva[1], Fabio Lapenta[1,2], Mateja Manček Keber[1], Roman Jerala[1,2,*], Mojca Benčina[1,2,*]

**Corresponding authors.** Roman Jerala,, +38614760335; Mojca Benčina,, +38614760334; (both) National Institute of Chemistry, Hajdrihova 19, SI-1001 Ljubljana, Slovenia, Fax: +38614760300.

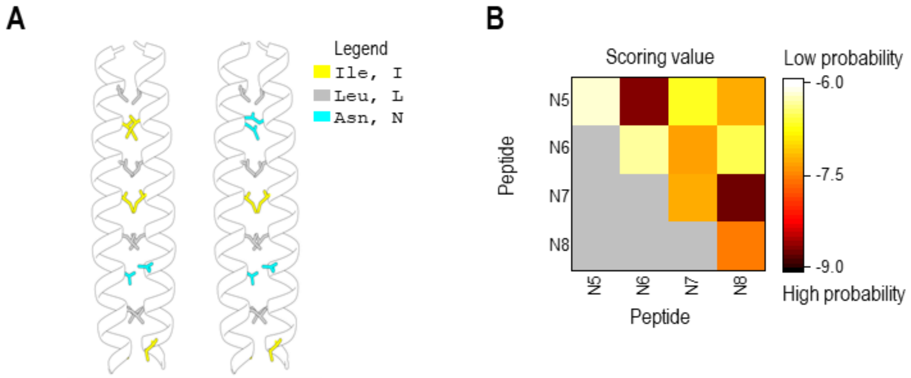


**Figure S1.** (**A**) Structural models of N7:N8 and P7A:P8A peptides built by the ISAMBARD modeling package. Selected amino acid side chains are shown for clarity. Polar amino acid residue Asn (cyan) is present at the *a* position of the second heptad of the N7, N8 (left). In P7A and P8A (right), additional Asn is placed at the *a* position of the fourth heptad. (**B**) Predicted orthogonality and interactions between all peptide pairs using a scoring algorithm[31]. The pairs, N5:N6 and N7:N8, were predicted to form CCs with higher stability.

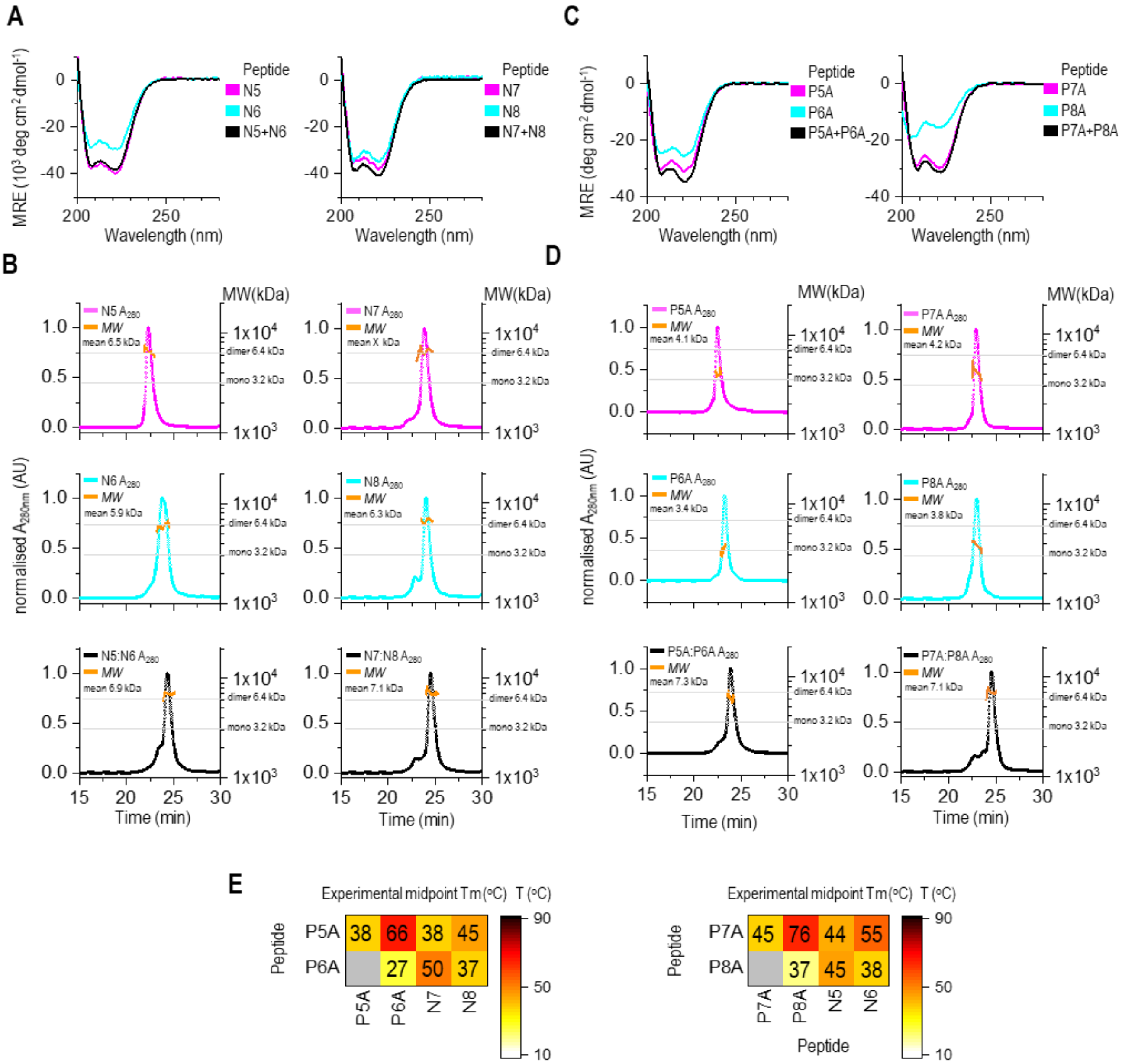


**Figure S2.** (**A**,**C**) Circular dichroism (CD) spectra of a 1:1 mixture of CC, N5:N6 and N7:N8 pairs, and P5A:P6A and P7A:P8A pairs (20 µM each, black). Single peptides (40 µM) are shown in cyan and magenta. The peptides and peptide mixtures resemble the characteristic α-helical spectrum at 20 ºC. The designated peptide partners exhibited a higher helical content than the peptides alone, indicating peptides' intrinsic preference for binding to their designated partners. All data were measured in Tris buffer. (**B**,**D**) Multimerization state of individual peptides and peptide pairs, N5:N6 and N7:N8, as determined by SEC-MALS. Size-exclusion chromatograms (normalized by setting the major peak maximum to 1) are presented with an overlay of the molecular weights (orange line) calculated from static light measurements. The dashed horizontal lines correspond to the expected molecular weights of a monomer and dimer. The SEC-MALS profiles of individual peptides confirm that the peptides P7A, P8A, P5A, and P6A are monomers in solution. (**E**) Heat map of the matrix of the calculated midpoint Tm from thermal denaturation scans of indicated peptide combinations.

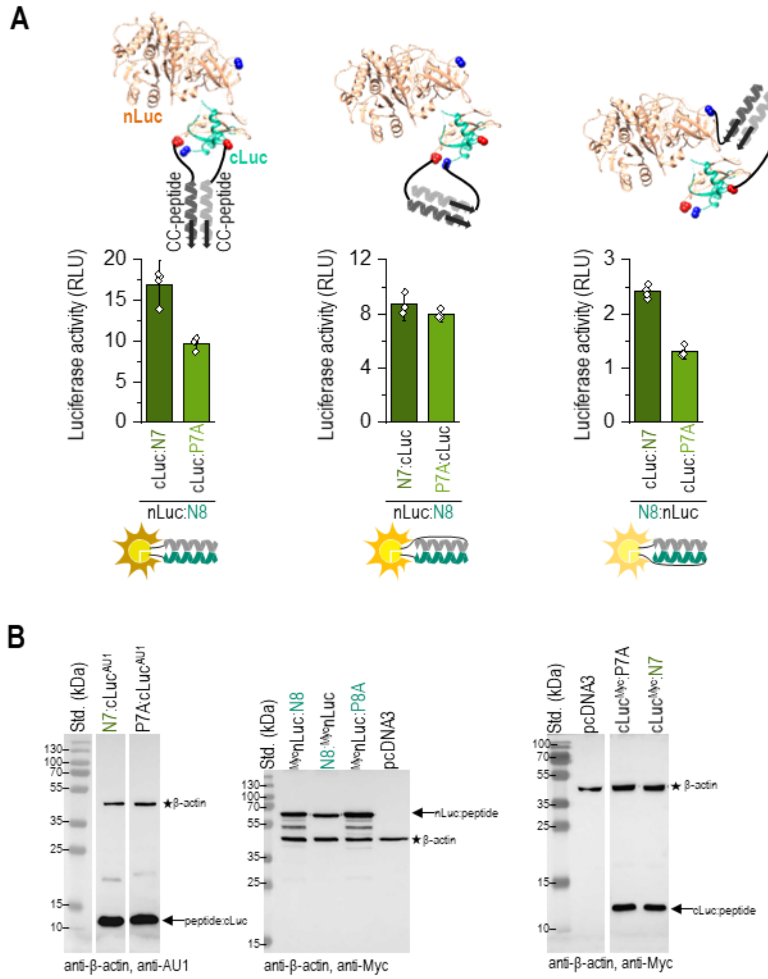


**Figure S3.** (**A**) Reconstitution of split luciferase in HEK293T cells. (Above) Schematic presentation of a structural model of split luciferase (brown nLuc and cyan cLuc) tethered to the N- or C-termini of the CC-forming peptide. *Note*: Linkers between CC peptides and N or C terminal of split luciferase have the same length. (Below) Luciferase activity of reconstituted CC-split luciferase 48 h after transfection of HEK293T cells with a plasmid expressing a combination of nLuc tethered to N8 and cLuc tethered to N7, or P7A peptides. The values represent the means (± s.d.) from four independent cell cultures and are representative of two independent experiments. Amounts of used plasmids are indicated in **Table S2**. (**B**) Expression of CC–split luciferase determined by a western blot test. Proteins separated via SDS-PAGE were blotted on a nitrocellulose membrane and stained with anti-Myc, anti-HA, and anti-β-actin antibodies as indicated.

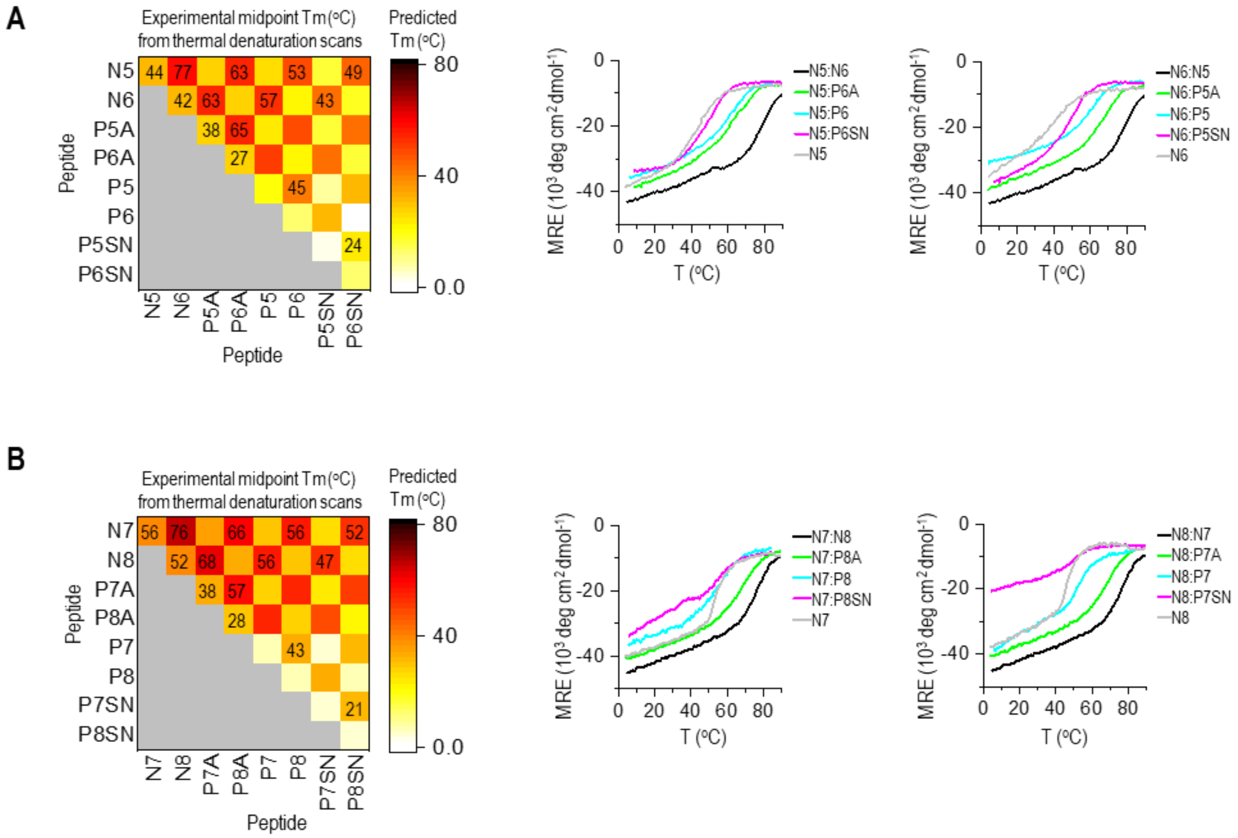


**Figure S4.** (**A, B**) The CC stability linear model [34] was used to predict the interactions between all possible peptide pairs. Predicted values are shown as a heat map. The experimental midpoint denaturation temperatures (Tm) of CCs and individual CC-forming peptides determined from thermal denaturation scans monitored by the CD signal at 222 nm (right) are depicted on top of the heat map of predicted Tm values.

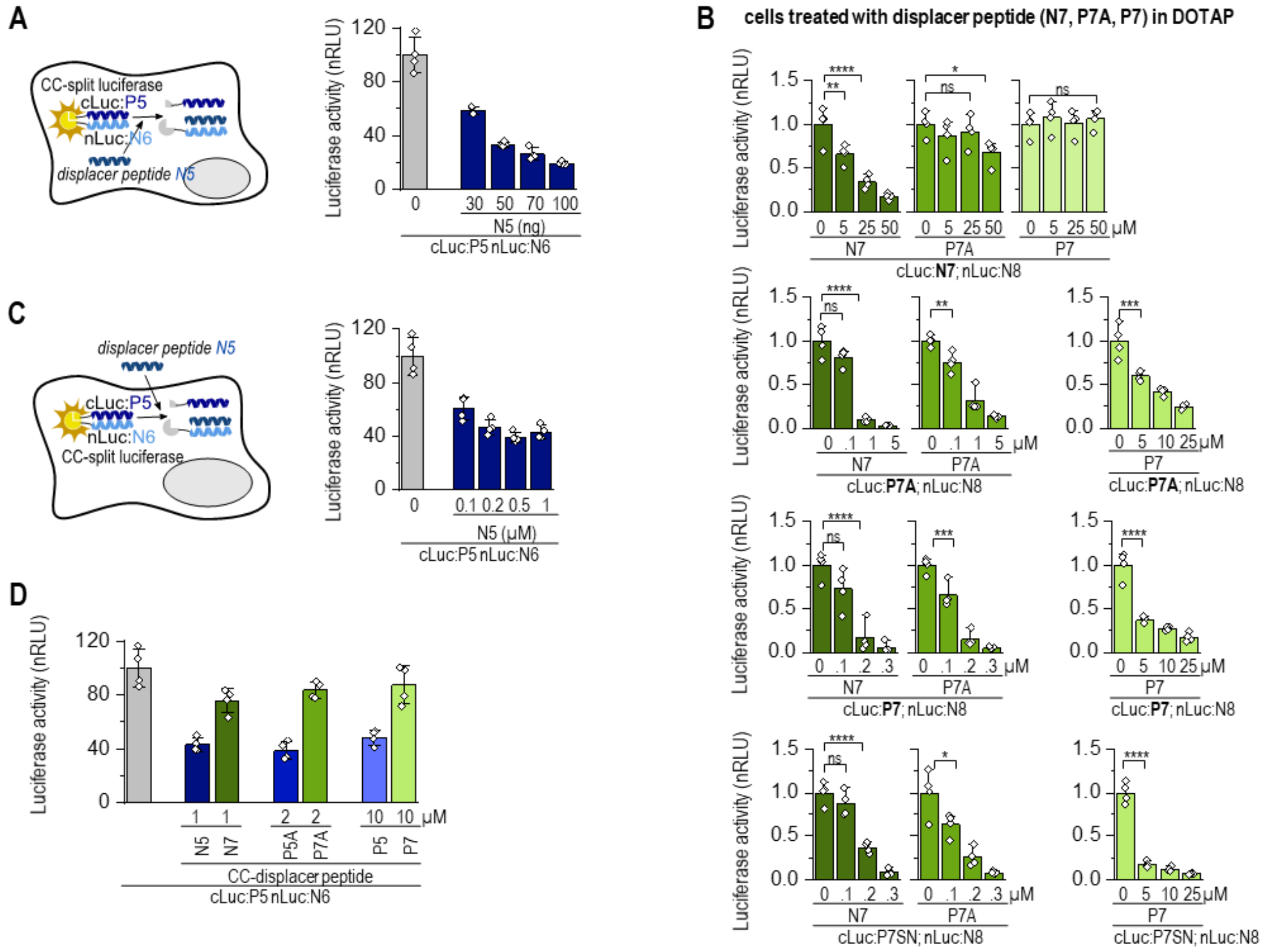


**Figure S5. (A)** Schematic representation of reconstituted N6:P5 split luciferase activity attenuated by N5 displacer peptide. Luciferase activity of HEK293T cells co-expressing cLuc:P5, nLuc:N6, and N5 displacer peptide. **(B)** Luciferase activity of HEK293T cells transfected with plasmids expressing nLuc:N8 and cLuc tethered with N7, P7A, P7, or P7SN. Forty-eight hours later, cells were treated with N7, P7A, or P7S displacer peptide (0–50 µM in DOTAP) for 2 h, and then luciferase activity was measured. **(C)** Schematic representation of reconstituted N6:P5 split luciferase activity attenuated by N5 displacer peptide, which was added to split luciferase-expressing cells. Luciferase activity of HEK293T cells treated with N5, P5A, P5, and P5SN peptide (0–1 µM in DOTAP) for 2 h. Cells were treated 48 h after transfection with plasmids expressing nLuc:N6 and cLuc tethered to P5. The bars represent the means (±s.d.) from four independent cell cultures. **(D)** Luciferase activity of HEK293T cells treated with N7 or N5; P7A or P5A; P7 or P5 peptide (1, 2, or 10 µM in DOTAP) for 2 h. Cells were treated 48 h after transfection with plasmids expressing nLuc:N6 and cLuc:P5. The values (**A-D**) represent the means (± s.d.) from four independent cell cultures, individually transfected with the same mixture of plasmids, and are representative of two independent experiments. For amounts of plasmids, see **Table S2**. Statistical analyses and the corresponding p-values are listed in **Table S3**.

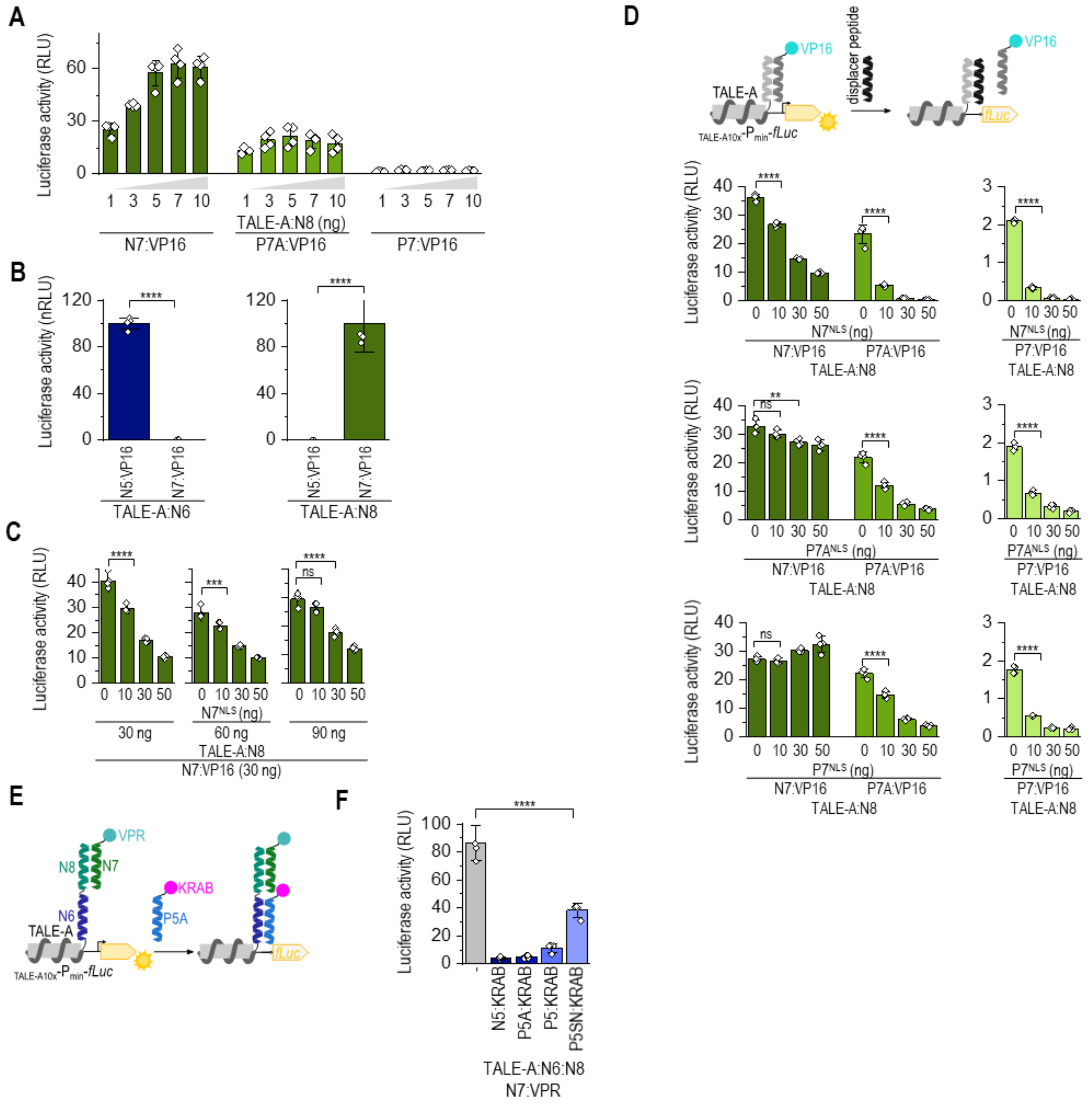


**Figure S6. Reconstitution of CC–split transcription factor attenuated with complementary CC–displacer peptide.** (**A**) Luciferase activity measured 48 h after transfection of HEK293T cells with plasmids expressing TALE-A:N8, CC:VP16 (30 ng) and the luciferase reporter. **(B)** Reconstitution of CC–split transcription factor in HEK293T cells co-expressing TALE-A:N8; VP16 linked to N7 or N5 or TALE-A:N6 and VP16 linked to N5 or N7; and the luciferase reporter ($_{TALE10x}$-P$_{min}$-fLuc). Luciferase activity was measured 48 h after transfection. (**C**) The amount of DNA-binding domain (TALE-A:N8) has a minor impact on displacement efficacy. Luciferase activity in HEK293T cells was measured 48 h after co-transfection of plasmids expressing TALE-A:N8, N7:VP16, N7$^{NLS}$ and reporter luciferase. (**D**) CC stability and CC–displacer peptide helicity determine displacement efficacy. Luciferase activity determined 48 h after transfection of HEK293T cells with plasmids expressing TALE:N8; CC:VP16; a CC–displacer peptide (N7$^{NLS}$, P7A$^{NLS}$ or P7$^{NLS}$); and the reporter luciferase ($_{TALE10x}$-P$_{min}$-fLuc). **(E)** Schematic representation of suppression of CC–split transcription factor activity (TALE:N6:N8-N7:VPR) by addition of CC:KRAB suppression domain. **(F)** Suppression of CC–split transcription factor activity (TALE:N6:N8-N7:VPR) by addition of KRAB suppression domain linked to N5, P5A, P5, or P5SN. Luciferase activity was measured 48 h after transfection. The values (**A-D,F**) represent the means (± s.d.) from four independent cell cultures, individually transfected with the same mixture of plasmids, and are representative of two independent experiments. For amounts of plasmids, see **Table S2**. Statistical analyses and the corresponding p-values are listed in **Table S3**.

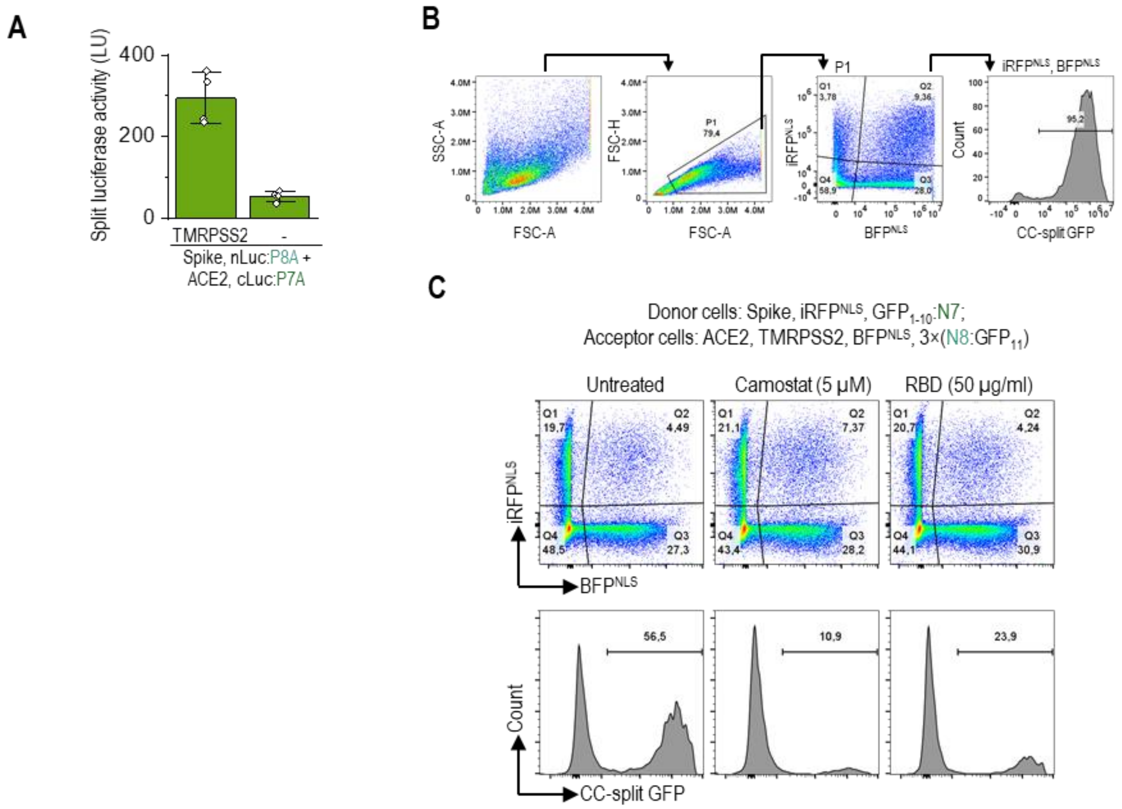


**Figure S7.** (**A**) Luciferase activity as an indicator of cell fusion. 24 h after transfection, donor HEK293T cells transfected with plasmids expressing nLuc:P8A, CoV-2 Spike-protein and acceptor cells expressing cLuc:P7A, ACE2, with or without TMPRSS2 were mixed in 1:1 ratio. Luciferase activity was measured 3 h later. The values represent the means (± s.d.) from four independent cell cultures, individually transfected with the same mixture of plasmids, and are representative of two independent experiments. (**B**) Flow cytometry gating strategy. The population of cells presented as pseudocolor plot (FCS-A/FCS-H) was gated for singlets and syncytia. The subset of cells, the BFP and iRFP positive, were analyzed for reconstituted CC-split GFP. The same gating strategy was used for **Fig 5E**, **Fig 6E,** and **Fig S7C**. (**C**) Flow cytometry analysis of a mixture of donor cells expressing the SARS CoV-2 Spike protein, $iRFP^{NLS}$, and $GFP_{1-10}$:N7 and the acceptor cells expressing ACE2 receptor, TMPRSS2, $BFP^{NLS}$, and 3×(N8:$GFP_{11}$). The populations of cells positive for iRFP and BFP are presented as a pseudocolor plot. Histograms present percent of reconstituted split GFP for double iRFP and BFP positive 3 h after mixing donor and acceptor cells. Formation of cell-cell fusion with split GFP reporter ($GFP_{1-10}$:N7; 3×(N8:$GFP_{11}$) analyzed by flow cytometry. Representative results of two independent experiments are shown. For amounts of plasmids, see **Table S2**.

**Table S1.** Sequences of orthogonal 5:6 and 7:8 N- and P-type peptides.

| Name | Sequence and heptad register gabcdef gabcdef gabcdef gabcdef | Electrostatic motif | Hydrophobic motif | Helicity* |
|---|---|---|---|---|
| **N7** | Y EIAALEA K**N**AALKA EIAALEA KIAALKA GC | EKEK | INII | 65 |
| **P7A** | YG EIAALEA K**N**AALKA EIAALEA K**N**AALKA GC | EKEK | ININ | 43 |
| P7 | SPED EIQALEE K**N**AQLKQ EIAALEE K**N**QALKY G | EKEK | ININ | 13 |
| P7SN | EIQQLEE K**N**SQLKQ EISQLEE K**N**QELKY G | EKEK | ININ | 3 |
| **N8** | Y KIAALKA E**N**AALEA KIAALKA EIAALEA GC | KEKE | INII | 59 |
| **P8A** | YG KIAALKA E**N**AALEA KIAALKA E**N**AALEA GGC | KEKE | ININ | 38 |
| P8 | SPED KIAQLKE E**N**QQLEQ KIQALKE E**N**AALEY G | KEKE | ININ | 11 |
| P8SN | KISELKE E**N**QQLEQ KIQQLKE E**N**SQLEY G | KEKE | ININ | 4 |
| **N5** | Y EIAALEA KIAALKA K**N**AALKA EIAALEA GC | EKKE | IINI | 61 |
| **P5A** | YG E**N**AALEA KIAALKA K**N**AALKA EIAALEA GC | EKKE | NINI | 51 |
| P5 | SPED E**N**AALEE KIAQLKQ K**N**AALKE EIQALEY G | EKKE | NINI | 20 |
| P5SN | E**N**SQLEE KISQLKQ K**N**SELKE EIQQLEY G | EKKE | NINI | 4 |
| **N6** | Y KIAALKA EIAALEA E**N**AALEA KIAALKA GC | KEEK | IINI | 60 |
| **P6A** | YG K**N**AALKA EIAALEA E**N**AALEA KIAALKA GGC | KEEK | NINI | 45 |
| P6 | SPED K**N**AALKE EIQALEE E**N**QALEE KIAQLKY G | KEEK | NINI | 16 |
| P6SN | K**N**SELKE EIQQLEE E**N**QQLEE KISELKY G | KEEK | NINI | 4 |

Legend: negative and positive amino acid residues are indicated with red and blue letters, respectively; asparagine is bolded;

* indicates that the value was calculated according to the method described by Agadir et al. [32].

**Table S2. Amounts of transfected plasmids for HEK293T cells in each well of a multi-well plate.** The empty pcDNA3 plasmid vector was used to equalize total DNA amounts up to 250 ng (for 96-well plate), 2500 ng (for 6-well plate), 1000 ng (for 8- and 12-well plate).

| Input plasmids | Amount [ng] |
|---|---|
| **Figure 1J (96-well plate)** | |
| cLuc:N7,<br>cluc:N5,<br>cLuc:P7A, or<br>cLuc:P5A | 30 |
| nLuc:N8, or<br>nLuc:N6 | 30 |
| phRL-TK | 5 |
| **Figure 1G (96-well plate)** | |
| cLuc:N7, or<br>N7:cLuc | 30 |
| nLuc:N8, or<br>N8:nLuc | 30 |
| phRL-TK | 5 |
| **Figures 1I, 1J (96-well plate)** | |
| cLuc:N7,<br>cLuc:P7A,<br>cLuc:N5, or<br>cLuc:P5A | 10-70 |
| nLuc:N8, or<br>nLuc:N6 | 30 |
| phRL-TK | 5 |
| **Figures 2A, 2B (96-well plate)** | |
| cLuc:N7,<br>cLuc:P7A,<br>cLuc:P7, or<br>cLuc:P7SNcLuc:N5,<br>cLuc:P5A,<br>cLuc:P5, or<br>cLuc:P5SN | 10-90 |
| nLuc:N8, or<br>nLuc:N6 | 30 |
| phRL-TK | 5 |
| **Figures 2C, 2D (96-well plate)** | |
| cLuc:N7,<br>cLuc:P7A,<br>cLuc:P7,<br>cLuc:P7SN,<br>cLuc:N5,<br>cLuc:P5A,<br>cLuc:P5, or<br>cLuc:P5SN | 10 |
| nLuc:N8, or<br>nLuc:N6 | 30 |
| phRL-TK | 5 |
| **Figure 3C (96-well plate)** | |
| cLuc:N7,<br>cLuc:P7A,<br>cLuc:P7, or<br>cLuc:P7SN | 30 |
| nLuc:N8 | 60 |
| N7 displacer peptide | 0-100 |
| phRL-TK | 50 |

| Input plasmids | Amount [ng] |
|---|---|
| **Figure 3E, S5B (96-well plate)** | |
| cLuc:N7,<br>cLuc:P7A,<br>cLuc:P7, or<br>cLuc:P7SN | 30 |
| nLuc:N8 | 60 |
| phRL-TK | 5 |
| **Figure 3G (96-well plate)** | |
| cLuc:P7 | 10 |
| nLuc:N8 | 30 |
| N7,<br>N5,<br>P7A,<br>P5A,<br>P7, or<br>P5 displacer peptide | 1-10 |
| phRL-TK | 5 |
| **Figure 4B (96-well plate)** | |
| TALA:NLS:N8, or<br>TALA:NLS:N6 | 30 |
| N7:NLS:VP16,<br>N5:NLS:VP6,<br>P7A:NLS:VP16 or<br>P5A:NLS:VP16 | 1-100 |
| phRL-TK | 5 |
| **Figure 4D (96-well plate)** | |
| TALA:NLS:N6:N8 | 30 |
| N7:VP16 | 30 |
| N5:KRAB,<br>P5A:KRAB,<br>P5:KRAB, or<br>P5SN:KRAB | 0-3 |
| phRL-TK | 5 |
| **Figure 4H (96-well plate)** | |
| TALA:NLS:N6:N8 | 30 |
| N7:NLS:VP16 | 30 |
| P5:KRAB | 1 |
| N5:$^{NLS}$ displacer peptide | 0-90 |
| phRL-TK | 5 |
| **Figure 4F (96-well plate)** | |
| TALA:NLS:N6:N8 | 30 |
| P7A:VP16 | 30 |
| N7$^{NLS}$ displacer peptide | 0-70 |
| phRL-TK | 5 |

| Input plasmids | Amount [ng] |
|---|---|
| **Figure 5B, 6B, 6D, S7A** | |
| **Acceptor cells (6-well plate)** | |
| cLuc:N7, or<br>cLuc:P7A | 1000 |
| ACE2 | 20 |
| TMPRSS2 | 0, 30 |
| **Donor cells (6-well plate)** | |
| nLuc:N8, or<br>nLuc:P8A | 1000 |
| S-protein | 10 |
| **Figure 5D, S7B, S7C** | |
| **Acceptor cells (8-well plate)** | |
| N8:$GFP_{11}$,<br>3x(N8:$GFP_{11}$), or<br>P8:$GFP_{11}$ | 100 |
| ACE2 | 40 |
| TMPRSS2 | 40 |
| BFP$^{NLS}$ | 120 |
| **Donor cells (8-well plate)** | |
| S-protein | 25 |
| $GFP_{1-10}$:N7, or<br>$GFP_{1-10}$:P7 | 100 |
| mCherry$^{NLS}$ | 500 |
| **Figure 5E, 6E(12-well plate)** | |
| **Acceptor cells (12-well plate)** | |
| N8:$GFP_{11}$,<br>3x(N8:$GFP_{11}$), or<br>P8:$GFP_{11}$ | 650 |
| ACE2 | 250 |
| TMPRSS2 | 50 |
| BFP$^{NLS}$ | 50 |
| **Donor cells (12-well plate)** | |
| S-protein | 50 |
| $GFP_{1-10}$:N7, or<br>$GFP_{1-10}$:P7, | 500 |
| iRFP$^{NLS}$ | 50 |

| Input plasmids | Amount [ng] |
|---|---|
| **Figure S3A (96-well plate)** | |
| cLuc:N7,<br>cluc:P7A,<br>N7:cLuc, or<br>P7A:cLuc | 30 |
| nLuc:N8, or<br>N8:nLuc | 60 |
| phRL-TK | 5 |
| **Figure S5A (96-well plate)** | |
| cLuc:P5 | 30 |
| N5 | 0-100 |
| nLuc:N6 | 30 |
| phRL-TK | 5 |
| **Figure S5C, S5D (96-well plate)** | |
| cLuc:P5 | 30 |
| nLuc:N6 | 30 |
| phRL-TK | 5 |
| **Figure S6A (96-well plate)** | |
| TALA:NLS:N8 | 1-10 |
| N7:NLS:VP16,<br>P7A:NLS:VP16 or<br>P7:NLS:VP16 | 30 |
| phRL-TK | 5 |
| **Figure S6B (96-well plate)** | |
| TALE-A:N8 or<br>TALE-A:N6 | 30 |
| N7:VP16 or<br>N5:VP16 | 30 |
| phRL-TK | 5 |
| **Figure S6c (96-well plate)** | |
| TALE-A:N8 | 30, 60, 80 |
| N7:VP16 | 30 |
| N7$^{NLS}$ | 0-50 |
| phRL-TK | 5 |
| **Figure S6D (96-well plate)** | |
| TALE-A:N8 | 30 |
| N7:VP16<br>P7A:VP16<br>P7:VP16 | 30 |
| N7$^{NLS}$<br>P7A$^{NLS}$<br>P7$^{NLS}$ | 0-50 |
| phRL-TK | 5 |
| **Figure S6F (96-well plate)** | |
| TALE-A:N6:N8 | 30 |
| N7:VPR | 60 |
| N5:KRAB<br>P5A:KRAB<br>P5:KRAB<br>P5SN:KRAB | 10 |
| phRL-TK | 5 |

### Table S3. Statistical analyses data.

| Figures | Test details | Significance | Summary | P-value |
|---|---|---|---|---|
| Fig 3C | **One-way ANOVA; Dunnett`s multiple comparisons test** | | | |
| | nLuc:N8 cLuc:N7; N7 | | | |
| | 0 *vs.* 80 | No | ns | 0.6788 |
| | 0 vs. 155 | No | ns | 0.2292 |
| | nLuc:N8 cLuc:P7A; N7 | | | |
| | 0 vs. 80 | Yes | **** | < 0,0001 |
| | 0 vs. 155 | Yes | **** | < 0,0001 |
| | nLuc:N8 cLuc:P7; N7 | | | |
| | 0 vs. 80 | Yes | **** | < 0,0001 |
| | 0 vs. 155 | Yes | **** | < 0,0001 |
| | nLuc:N8 cLuc:P7SN; N7 | | | |
| | 0 vs. 80 | Yes | **** | < 0,0001 |
| | 0 vs. 155 | Yes | **** | < 0,0001 |
| Fig 3E | **One-way ANOVA; Dunnett`s multiple comparisons test** | | | |
| | cLuc:N7 nLuc:N8; N7 | | | |
| | 0 vs. 0.1 | No | ns | 0.9861 |
| | 0 vs. 0.2 | No | ns | 0.9997 |
| | 0 vs. 0.4 | No | ns | 0.8104 |
| | 0 vs. 1 | No | ns | > 0,9999 |
| | 0 vs. 2 | No | ns | 0.9997 |
| | 0 vs. 5 | No | ns | 0.5602 |
| | 0 vs. 10 | Yes | ** | 0.0037 |
| | **One-way ANOVA Dunnett`s multiple comparisons test** | | | |
| | cLuc:P7A nLuc:N8; N7 | | | |
| | 0 *vs.* n (for all tested amounts) | Yes | **** | < 0,0001 |
| | cLuc:P7 nLuc:N8; N7 | | | |
| | 0 *vs.* n (for all tested amounts) | Yes | **** | < 0,0001 |
| | cLuc:P7SN nLuc:N8; N7 | | | |
| | 0 *vs.* n (for all tested amounts) | Yes | **** | < 0,0001 |
| Fig S5B | **One-way ANOVA; Dunnett`s multiple comparisons test** | | | |
| | nLuc:N8 cLuc:N7; N7 | | | |
| | 0 *vs.* 5 | Yes | ** | 0.0072 |
| | 0 *vs.* 25 | Yes | **** | < 0,0001 |
| | 0 *vs.* 50 | Yes | **** | < 0,0001 |
| | nLuc:N8 cLuc:N7; P7A | | | |
| | 0 *vs.* 5 | No | ns | 0.5576 |
| | 0 *vs.* 25 | No | ns | 0.8284 |
| | 0 *vs.* 50 | Yes | * | 0.0445 |
| | nLuc:N8 cLuc:N7; P7 | | | |
| | 0 *vs.* 5 | No | ns | 0.7673 |
| | 0 *vs.* 25 | No | ns | 0.9968 |
| | 0 *vs.* 50 | No | ns | 0.8588 |
| | nLuc:N8 cLuc:P7A; N7 | | | |
| | 0 *vs.* 0.1 | No | ns | 0.0575 |
| | 0 *vs.* 1 | Yes | **** | < 0,0001 |
| | 0 *vs.* 5 | Yes | **** | < 0,0001 |
| | nLuc:N8 cLuc: P7A; P7A | | | |
| | 0 *vs.* 0.1 | Yes | ** | 0.009 |
| | 0 *vs.* 1 | Yes | **** | < 0,0001 |
| | 0 *vs.* 5 | Yes | **** | < 0,0001 |
| | nLuc:N8 cLuc: P7A; P7 | | | |
| | 0 *vs.* 0.1 | Yes | *** | 0.0003 |
| | 0 *vs.* 1 | Yes | **** | < 0,0001 |
| | 0 *vs.* 5 | Yes | **** | < 0,0001 |
| | nLuc:N8 cLuc:P7; N7 | | | |
| | 0 *vs.* 0.1 | No | ns | 0.1021 |
| | 0 *vs.* 0.2 | Yes | **** | < 0,0001 |
| | 0 *vs.* 0.3 | Yes | **** | < 0,0001 |
| | nLuc:N8 cLuc: P7; P7A | | | |
| | 0 *vs.* 0.1 | Yes | *** | 0.0006 |
| | 0 *vs.* 0.2 | Yes | **** | < 0,0001 |
| | 0 *vs.* 0.3 | Yes | **** | < 0,0001 |
| | nLuc:N8 cLuc: P7; P7 | | | |
| | 0 *vs.* n (for all tested amounts) | Yes | **** | < 0,0001 |
| | nLuc:N8 cLuc:P7SN; N7 | | | |
| | 0 *vs.* 0.1 | No | ns | 0.276 |
| | 0 *vs.* 0.2 | Yes | **** | < 0,0001 |
| | 0 *vs.* 0.3 | Yes | **** | < 0,0001 |
| | nLuc:N8 cLuc: P7SN; P7A | | | |
| | 0 *vs.* 0.1 | Yes | * | 0.0167 |
| | 0 *vs.* 0.2 | Yes | **** | < 0,0001 |
| | 0 *vs.* 0.3 | Yes | **** | < 0,0001 |
| | nLuc:N8 cLuc: P7SN; P7 | | | |
| | 0 *vs.* n (for all tested amounts) | Yes | **** | < 0,0001 |

| Figures | Test details | Significance | Summary | P-value |
|---|---|---|---|---|
| Fig 4D | **One-way ANOVA; Dunnett`s multiple comparisons test** | | | |
| | TALE-A:N6:N8 N7:VP16, N5:KRAB | | | |
| | 0 *vs.* n (for all tested amounts) | Yes | **** | < 0,0001 |
| | TALE-A:N6:N8 N7:VP16, P5A:KRAB | | | |
| | 0 *vs.* n (for all tested amounts) | Yes | **** | < 0,0001 |
| | TALE-A:N6:N8 N7:VP16, P5:KRAB | | | |
| | 0 *vs.* n (for all tested amounts) | Yes | ** | 0.001 |
| | TALE-A:N6:N8 N7:VP16, P5SN:KRAB | | | |
| | 0 *vs.* n (for all tested amounts) | Yes | **** | < 0,0001 |
| Fig 4F | **One-way ANOVA; Dunnett`s multiple comparisons test** | | | |
| | TALE-A:N6:N8 P7A:VP16, N7$^{NLS}$:KRAB | | | |
| | 0 *vs.* n (for all tested amounts) | Yes | **** | < 0,0001 |
| Fig 4H | **One-way ANOVA; Dunnett`s multiple comparisons test** | | | |
| | - *vs.* 0 | Yes | **** | < 0,0001 |
| | - *vs.* 10 | Yes | **** | < 0,0001 |
| | - *vs.* 30 | No | ns | 0.2404 |
| | - *vs.* 50 | No | ns | 0.2701 |
| | - *vs.* 70 | Yes | *** | 0.0004 |
| Fig S6B | **Paired t-test, two-tailed** | | | |
| | TALE-A:N6 N5:VP16 vs. N7:VP16 | Yes | **** | < 0,0001 |
| | TALE-A:N6 N8:VP16 vs. N7:VP16 | Yes | **** | < 0,0001 |
| Fig S6C | **One-way ANOVA; Dunnett`s multiple comparisons test** | | | |
| | TALE-A:N8 N7:VP16, N7$^{NLS}$ | | | |
| | 0 *vs.* n (for all tested amounts) | Yes | **** | < 0,0001 |
| | TALE-A:N8 N7:VP16, N7$^{NLS}$ | | | |
| | 0 *vs.* n (for all tested amounts) | Yes | **** | < 0,0001 |
| | TALE-A:N8 N7:VP16, N7$^{NLS}$ | | | |
| | 0 *vs.* 10 | No | ns | 0.0638 |
| | 0 *vs.* 30 | Yes | **** | < 0,0001 |
| | 0 *vs.* 50 | Yes | **** | < 0,0001 |
| Fig S6D | **One-way ANOVA; Dunnett`s multiple comparisons test** | | | |
| | TALE-A:N8 N7:VP16, N7$^{NLS}$ | | | |
| | 0 *vs.* n (for all tested amounts) | Yes | **** | < 0,0001 |
| | TALE-A:N8 P7A:VP16, N7$^{NLS}$ | | | |
| | 0 *vs.* n (for all tested amounts) | Yes | **** | < 0,0001 |
| | TALE-A:N8 P7:VP16, N7$^{NLS}$ | | | |
| | 0 *vs.* n (for all tested amounts) | Yes | **** | < 0,0001 |
| | TALE-A:N8 N7:VP16, P7A$^{NLS}$ | | | |
| | 0 *vs.* 10 | No | ns | 0.086 |
| | 0 *vs.* 30 | Yes | ** | 0.0011 |
| | 0 *vs.* 50 | Yes | *** | 0.0002 |
| | TALE-A:N8 P7A:VP16, P7A$^{NLS}$ | | | |
| | 0 *vs.* n (for all tested amounts) | Yes | **** | < 0,0001 |
| | TALE-A:N8 P7:VP16, P7A$^{NLS}$ | | | |
| | 0 *vs.* n (for all tested amounts) | Yes | **** | < 0,0001 |
| | TALE-A:N8 N7:VP16, P7$^{NLS}$ | | | |
| | 0 *vs.* 10 | No | ns | 0.9364 |
| | 0 *vs.* 30 | No | ns | 0.0507 |
| | 0 *vs.* 50 | Yes | ** | 0.0018 |
| | TALE-A:N8 P7A:VP16, P7$^{NLS}$ | | | |
| | 0 *vs.* n (for all tested amounts) | Yes | **** | < 0,0001 |
| | TALE-A:N8 P7:VP16, P7$^{NLS}$ | | | |
| | 0 *vs.* n (for all tested amounts) | Yes | **** | < 0,0001 |
| Fig S6F | **One-way ANOVA; Dunnett`s multiple comparisons test** | | | |
| | TALE-A:N8 N7:VPR | | | |
| | - *vs.* N5:KRAB | Yes | **** | < 0,0001 |
| | - *vs.* P5A:KRAB | Yes | **** | < 0,0001 |
| | - *vs.* P5:KRAB | Yes | **** | < 0,0001 |
| | - *vs.* P5SN:KRAB | Yes | **** | < 0,0001 |
| Fig 5B | **Paired t-test, two-tailed** | | | |
| | nLuc:N8 cLuc:N7 vs. nLuc:P8A cLuc:P7A | No | ns | 0.8335 |
| Fig 6B | **One-way ANOVA; Dunnett`s multiple comparisons test** | | | |
| | Spike nLuc:P8A, TMRSS2 ACE2 cLuc:P7A, Camostat | | | |
| | 0 *vs.* n (for all tested amounts) | Yes | **** | < 0,0001 |
| Fig 6D | **One-way ANOVA; Dunnett`s multiple comparisons test** | | | |
| | Spike nLuc:P8A, TMRSS2 ACE2 cLuc:P7A, RBD | | | |
| | 0 *vs.* n (for all tested amounts) | Yes | **** | < 0,0001 |

**Table S4.** The list of plasmids used in this study.

| **Plasmid name (amino acid sequence)** |
|---|
| Myc:N7 |
| M EQKLISEEDL GEIAALEAKNAALKAEIAALEAKIAALKAGY* |
| red: Myc tag; black: N7 |
| Myc:P7A |
| M EQKLISEEDL GEIAALEAKNAALKAEIAALEAKNAALKAGC* |
| red: Myc tag; black: P7 |
| Myc:P7 |
| M EQKLISEEDL EIQALEEKNAQLKQEIAALEEKNQALKYG* |
| red: Myc tag; black: P7 |
| Myc:P7SN |
| M EQKLISEEDL EIQQLEEKNSQLKQEISQLEEKNQELKYG* |
| red: Myc tag; black: P7SN |
| Myc:N7Q |
| M EQKLISEEDL GEIAALEQKNAALKQEIAALEQKIAALKQGC* |
| red: Myc tag; black: N7Q |
| N7:gs linker:cLuc:gs linker:HA |
| M GEIAALEAKNAALKAEIAALEAKIAALKAGY GGSGGGGSGGS TMTEKEIVDYVASQVTTAKKLRGGVVFVDEVPKGLTGKLDARKIREILIKAKKGGKIAVNS GSG<br>YPYDVPDYA* |
| black: N7; dark blue: gs linker; yellow: C terminal of split Luciferase cLuc; green:HA tag |
| N7Q:gs linker:cLuc:gs linker: HA |
| M GEIAALEQKNAALKQEIAALEQKIAALKQGC GGSGGGSGGS TMTEKEIVDYVASQVTTAKKLRGGVVFVDEVPKGLTGKLDARKIREILIKAKKGGKIAVNS GSG<br>YPYDVPDYA* |
| black: N7Q; dark blue: gs linker; yellow: C terminal of split Luciferase cLuc; green:HA tag |
| P7A:gs linker:cLuc:gs linker:HA |
| M GEIAALEAKNAALKAEIAALEAKNAALKAGC GGSGGGGSGG TMTEKEIVDYVASQVTTAKKLRGGVVFVDEVPKGLTGKLDARKIREILIKAKKGGKIAVN SGSG<br>YPYDVPDYA* |
| black: P7A; dark blue: gs linker; yellow: C terminal of split Luciferase cLuc; green:HA tag |
| P7:gs linker:cLuc:gs linker:HA |
| M EIQALEEKNAQLKQEIAALEEKNQALKYG GGSGGGGSGGS TMTEKEIVDYVASQVTTAKKLRGGVVFVDEVPKGLTGKLDARKIREILIKAKKGGKIAVN SGSG<br>YPYDVPDYA* |
| black: P7; dark blue: gs linker; yellow: C terminal of split Luciferase cLuc; green:HA tag |
| P7SN:gs linker:cLuc:gs linker:HA |
| M EIQQLEEKNSQLKQEISQLEEKNQELKYG GGSGGGGSGGS TMTEKEIVDYVASQVTTAKKLRGGVVFVDEVPKGLTGKLDARKIREILIKAKKGGKIAVN SGSG<br>YPYDVPDYA* |
| black: P7SN; dark blue: gs linker; yellow: C terminal of split Luciferase cLuc; green:HA tag |
| cLuc:Myc:N7 |
| M TMTEKEIVDYVASQVTTAKKLRGGVVFVDEVPKGLTGKLDARKIREILIKAKKGGKIAVNS EQKLISEEDL GEIAALEAKNAALKAEIAALEAKIAALKAGY |
| yellow: C terminal of split Luciferase cLuc; red:Myc tag; dark blue: gs linker; black: N7 |
| cLuc:Myc: N7Q |
| M TMTEKEIVDYVASQVTTAKKLRGGVVFVDEVPKGLTGKLDARKIREILIKAKKGGKIAVNS EQKLISEEDL GEIAALEQKNAALKQEIAALEQKIAALKQGC |
| yellow: C terminal of split Luciferase cLuc; red:Myc tag; dark blue: gs linker; black: N7 |
| cLuc:Myc: P7A |
| M TMTEKEIVDYVASQVTTAKKLRGGVVFVDEVPKGLTGKLDARKIREILIKAKKGGKIAVN EQKLISEEDL GEIAALEAKNAALKAEIAALEAKNAALKAGC |
| yellow: C terminal of split Luciferase cLuc; red:Myc tag; dark blue: gs linker; black: N7 |
| cLuc:Myc: P7 |
| M TMTEKEIVDYVASQVTTAKKLRGGVVFVDEVPKGLTGKLDARKIREILIKAKKGGKIAVN EQKLISEEDL EIQALEEKNAQLKQEIAALEEKNQALKYG |
| yellow: C terminal of split Luciferase cLuc; red:Myc tag; dark blue: gs linker; black: N7 |
| cLuc:Myc: P7SN |
| M TMTEKEIVDYVASQVTTAKKLRGGVVFVDEVPKGLTGKLDARKIREILIKAKKGGKIAVN EQKLISEEDL EIQQLEEKNSQLKQEISQLEEKNQELKYG |
| yellow: C terminal of split Luciferase cLuc; red:Myc tag; dark blue: gs linker; black: N7 |
| nLuc:myc:gs linker:N8 |
| MGSGEDAKNIKKGPAPFYPLEDGTAGEQLHKAMKRYALVPGTIAFTDAHIEVDITYAEYFEMSVRLAEAMKRYGLNTNHRIVVCSENSLQFFMPVLGALFIGVAVAPAND<br>IYNERELLNSMGISQPTVVFVSKKGLQKILNVQKKLPIIQKIIIMDSKTDYQGFQSMYTFVTSHLPPGFNEYDFVPESFDRDKTIALIMNSSGSTGLPKGVALPHRTACV<br>RFSHARDPIFGNQIIPDTAILSVVPFHHGFGMFTTLGYLICGFRVVLMYRFEEELFLRSLQDYKIQSALLVPTLFSFFAKSTLIDKYDLSNLHEIASGGAPLSKEVGEAV<br>AKRFHLPGIRQGYGLTETTSAILITPEGDDKPGAVGKVVPFFEAKVVDLDTGKTLGVNQRGELCVRGPMIMSGYVNNPEATNALIDKDGWLHSGDIAYWDEDEHFFIVDR<br>LKSLIKYKGYQVAPAELESILLQHPNIFDAGVAGLPDDDAGELPAAVVVLEHGK EQKLISEEDL GGSGG YGKIAALKAENAALEAKIAALKAEIAALEAGY* |
| purple: N terminal of split luciferase nLuc; red: Myc tag; dark blue: gs linker; black: N8 |
| N8:myc: gs linker:nLuc |
| M YGKIAALKAENAALEAKIAALKAEIAALEAGY EQKLISEEDL GGSGGGSG<br>EDAKNIKKGPAPFYPLEDGTAGEQLHKAMKRYALVPGTIAFTDAHIEVDITYAEYFEMSVRLAEAMKRYGLNTNHRIVVCSENSLQFFMPVLGALFIGVAVAPANDIYNE<br>RELLNSMGISQPTVVFVSKKGLQKILNVQKKLPIIQKIIIMDSKTDYQGFQSMYTFVTSHLPPGFNEYDFVPESFDRDKTIALIMNSSGSTGLPKGVALPHRTACVRFSH<br>ARDPIFGNQIIPDTAILSVVPFHHGFGMFTTLGYLICGFRVVLMYRFEEELFLRSLQDYKIQSALLVPTLFSFFAKSTLIDKYDLSNLHEIASGGAPLSKEVGEAVAKRF<br>HLPGIRQGYGLTETTSAILITPEGDDKPGAVGKVVPFFEAKVVDLDTGKTLGVNQRGELCVRGPMIMSGYVNNPEATNALIDKDGWLHSGDIAYWDEDEHFFIVDRLKSL<br>IKYKGYQVAPAELESILLQHPNIFDAGVAGLPDDDAGELPAAVVVLEHGK* |
| purple: N terminal of split luciferase nLuc; red: Myc tag; dark blue: gs linker; black: N8 |
| His:TALE-A:NLS:gs lineker:N8 |
| M HHHHHH<br>DYKDHDGDYKDHDIDYKDDDDKMAPKKKRKVGIHRGVPMVDLRTLGYSQQQQEKIKPKVRSTVAQHHEALVGHGFTHAHIVALSQHPAALGTVAVKYQDMIAALPEATHE<br>AIVGVGKQWSGARALEALLTVAGELRGPPLQLDTGQLLKIAKRGGVTAVEAVHAWRNALTGAPLNLTPDQVVAIASNGGGKQALETVQRLLPVLCQDHGLTPEQVVAIAS<br>NGGGKQALETVQRLLPVLCQAHGLTPDQVVAIASNIGGKQALETVQRLLPVLCQAHGLTPAQVVAIASHDGGKQALETVQRLLPVLCQDHGLTPDQVVAIASNGGGKQAL |

ETVQRLLPVLCQDHGLTPEQVVAIANNNGGKQALETVQRLLPVLCQAHGLTPDQVVAIASHDGGKQALETVQRLLPVLCQAHGLTPAQVVAIASNGGGKQALETVQRLLP
VLCQDHGLTPDQVVAIANNNGGKQALETVQRLLPVLCQDHGLTPEQVVAIASHDGGKQALETVQRLLPVLCQAHGLTPDQVVAIASNGGGKQALETVQRLLPVLCQAHGL
TPAQVVAIASHDGGKQALETVQRLLPVLCQDHGLTPDQVVAIASHDGGKQALETVQRLLPVLCQDHGLTPEQVVAIASHDGGKQALETVQRLLPVLCQAHGLTPDQVVAI
ANNNGGKQALETVQRLLPVLCQAHGLTPAQVVAIASHDGGKQALETVQRLLPVLCQDHGLTPEQVVAIASNGGGRPALESIVAQLSRPDPALAALTNDHLVALACLGGRP
ALDAVKKGLPHAPALIKRTNRRIPERTSHRVAGS DPKKKRKV GGSGGGSGGS YKIAALKAENAALEAKIAALKAEIAALEAGC*

magenta: his tag; orange: TALA; dark blue: gs linker; cian: nuclear localization sequence; black: N8

### N7:gs linker:NLS:VP16:[AU1]

M GEIAALEAKNAALKAEIAALEAKIAALKAGY GGSGGGSGGS DPKKKRKV
APPTDVSLGDELHLDGEDVAMAHADALDDFDLDMLGDGDSPGPGFTPHDSAPYGALDMADFEFEQMFTDALGIDEYGG DTYRYI*

black:N7; dark blue:gs linker; cian: nuclear localization sequence; dark green VP16 activation domain; light green: AU1 tag

### N7Q:gs linker:NLS:VP16:[AU1]

M GEIAALEQKNAALKQEIAALEQKIAALKQGC GGSGGGSGGS DPKKKRKV
APPTDVSLGDELHLDGEDVAMAHADALDDFDLDMLGDGDSPGPGFTPHDSAPYGALDMADFEFEQMFTDALGIDEYGG DTYRYI*

black:N7Q; dark blue:gs linker; cian: nuclear localization sequence; dark green VP16 activation domain; light green: AU1 tag

### P7A:gs linker:NLS:VP16:[AU1]

M GEIAALEAKNAALKAEIAALEAKNAALKAGC GGSGGGSGGS DPKKKRKV
APPTDVSLGDELHLDGEDVAMAHADALDDFDLDMLGDGDSPGPGFTPHDSAPYGALDMADFEFEQMFTDALGIDEYGG DTYRYI*

black:P7A; dark blue:gs linker; cian: nuclear localization sequence; dark green VP16 activation domain; light green: AU1 tag

### P7:gs linker:NLS:VP16:[AU1]

M EIQALEEKNAQLKQEIAALEEKNQALKYG GGSGGGSGGS DPKKKRKV
APPTDVSLGDELHLDGEDVAMAHADALDDFDLDMLGDGDSPGPGFTPHDSAPYGALDMADFEFEQMFTDALGIDEYGG DTYRYI*

black:P7; dark blue:gs linker; cian: nuclear localization sequence; dark green VP16 activation domain; light green: AU1 tag

### P7SN:gs linker:NLS:VP16:[AU1]

M EIQQLEEKNSQLKQEISQLEEKNQELKY GGSGGGSGGS DPKKKRKV
APPTDVSLGDELHLDGEDVAMAHADALDDFDLDMLGDGDSPGPGFTPHDSAPYGALDMADFEFEQMFTDALGIDEYGG DTYRYI*

black:P7SN; dark blue:gs linker; cian: nuclear localization sequence; dark green VP16 activation domain; light green: AU1 tag

### Myc:N7:NLS

M EQKLISEEDL GEIAALEAKNAALKAEIAALEAKIAALKAGY DPKKKRKV*

red: Myc tag; Black: N7; cian: nuclear localization sequence

### Myc:N7Q:NLS

M EQKLISEEDL GEIAALEQKNAALKQEIAALEQKIAALKQGC DPKKKRKV*

red: Myc tag; Black: N7Q; cian: nuclear localization sequence

### Myc:P7A:NLS

M EQKLISEEDL GEIAALEAKNAALKAEIAALEAKNAALKAGC DPKKKRKV*

red: Myc tag; Black: P7A; cian: nuclear localization sequence

### Myc:P7:NLS

M EQKLISEEDL EIQALEEKNAQLKQEIAALEEKNQALKYG DPKKKRKV*

red: Myc tag; Black: P7; cian: nuclear localization sequence

### Myc:P7SN:NLS

M EQKLISEEDL EIQQLEEKNSQLKQEISQLEEKNQELKYG DPKKKRKV*

red: Myc tag; Black: P7SN; cian: nuclear localization sequence

### nLuc:P8A

MGSGEDAKNIKKGPAPFYPLEDGTAGEQLHKAMKRYALVPGTIAFTDAHIEVDITYAEYFEMSVRLAEAMKRYGLNTNHRIVVCSENSLQFFMPVLGALFIGVAVAPAND
IYNERELLNSMGISQPTVVFVSKKGLQKILNVQKKLPIIQKIIIMDSKTDYQGFQSMYTFVTSHLPPGFNEYDFVPESFDRDKTIALIMNSSGSTGLPKGVALPHRTACV
RFSHARDPIFGNQIIPDTAILSVVPFHHGFGMFTTLGYLICGFRVVLMYRFEEELFLRSLQDYKIQSALLVPTLFSFFAKSTLIDKYDLSNLHEIASGGAPLSKEVGEAV
AKRFHLPGIRQGYGLTETTSAILITPEGDDKPGAVGKVVPFFEAKVVDLDTGKTLGVNQRGELCVRGPMIMSGYVNNPEATNALIDKDGWLHSGDIAYWDEDEHFFIVDR
LKSLIKYKGYQVAPAELESILLQHPNIFDAGVAGLPDDDAGELPAAVVVLEHGK EQKLISEEDL GGSGG KIAALKAENAALEAKIAALKAENAALEAGGC

purple: N terminal of split luciferase nLuc; red: Myc tag; dark blue: gs linker; black: P8A

### cLuc:N5

M TMTEKEIVDYVASQVTTAKKLRGGVVFVDEVPKGLTGKLDARKIREILIKAKKGGKIAVN EQKLISEEDL GEIAALEAKIAALKAKNAALKAEIAALEAGY

yellow: C terminal of split Luciferase cLuc; red:Myc tag; dark blue: gs linker; black: N5

### cLuc:P5A

M TMTEKEIVDYVASQVTTAKKLRGGVVFVDEVPKGLTGKLDARKIREILIKAKKGGKIAVN EQKLISEEDL ENAALEAKIAALKAKNAALKAEIAALEA

yellow: C terminal of split Luciferase cLuc; red:Myc tag; dark blue: gs linker; black: P5A

### cLuc:P5

M TMTEKEIVDYVASQVTTAKKLRGGVVFVDEVPKGLTGKLDARKIREILIKAKKGGKIAVN EQKLISEEDL SPEDENAALEEKIAQLKQKNAALKEEIQALEYG

yellow: C terminal of split Luciferase cLuc; red:Myc tag; dark blue: gs linker; black: P5

### cLuc:myc:P5SN

M TMTEKEIVDYVASQVTTAKKLRGGVVFVDEVPKGLTGKLDARKIREILIKAKKGGKIAVN EQKLISEEDL ENSQLEEKISQLKQKNSELKEEIQQLE

yellow: C terminal of split Luciferase cLuc; red:Myc tag; dark blue: gs linker; black: P5SN

### nLuc:P6A

MGSGEDAKNIKKGPAPFYPLEDGTAGEQLHKAMKRYALVPGTIAFTDAHIEVDITYAEYFEMSVRLAEAMKRYGLNTNHRIVVCSENSLQFFMPVLGALFIGVAVAPAND
IYNERELLNSMGISQPTVVFVSKKGLQKILNVQKKLPIIQKIIIMDSKTDYQGFQSMYTFVTSHLPPGFNEYDFVPESFDRDKTIALIMNSSGSTGLPKGVALPHRTACV
RFSHARDPIFGNQIIPDTAILSVVPFHHGFGMFTTLGYLICGFRVVLMYRFEEELFLRSLQDYKIQSALLVPTLFSFFAKSTLIDKYDLSNLHEIASGGAPLSKEVGEAV
AKRFHLPGIRQGYGLTETTSAILITPEGDDKPGAVGKVVPFFEAKVVDLDTGKTLGVNQRGELCVRGPMIMSGYVNNPEATNALIDKDGWLHSGDIAYWDEDEHFFIVDR
LKSLIKYKGYQVAPAELESILLQHPNIFDAGVAGLPDDDAGELPAAVVVLEHGK EQKLISEEDL GGSGG GKNAALKAEIAALEAENAALEAKIAALKAGGC

purple: N terminal of split luciferase nLuc; red: Myc tag; dark blue: gs linker; black:P6A

### nLuc:N6

MGSGEDAKNIKKGPAPFYPLEDGTAGEQLHKAMKRYALVPGTIAFTDAHIEVDITYAEYFEMSVRLAEAMKRYGLNTNHRIVVCSENSLQFFMPVLGALFIGVAVAPAND
IYNERELLNSMGISQPTVVFVSKKGLQKILNVQKKLPIIQKIIIMDSKTDYQGFQSMYTFVTSHLPPGFNEYDFVPESFDRDKTIALIMNSSGSTGLPKGVALPHRTACV
RFSHARDPIFGNQIIPDTAILSVVPFHHGFGMFTTLGYLICGFRVVLMYRFEEELFLRSLQDYKIQSALLVPTLFSFFAKSTLIDKYDLSNLHEIASGGAPLSKEVGEAV
AKRFHLPGIRQGYGLTETTSAILITPEGDDKPGAVGKVVPFFEAKVVDLDTGKTLGVNQRGELCVRGPMIMSGYVNNPEATNALIDKDGWLHSGDIAYWDEDEHFFIVDR
LKSLIKYKGYQVAPAELESILLQHPNIFDAGVAGLPDDDAGELPAAVVVLEHGK EQKLISEEDL GGSGG KIAALKAEIAALEAENAALEAKIAALKAG*

purple: N terminal of split luciferase nLuc; red: Myc tag; dark blue: gs linker; black:N6

### N5:gs linker:NLS:VP16:[AU1]

M GEIAALEAKIAALKAKNAALKAEIAALEAGY GGSGGGGSGGS DPKKKRKV
APPTDVSLGDELHLDGEDVAMAHADALDDFDLDMLGDGDSPGPGFTPHDSAPYGALDMADFEFEQMFTDALGIDEYGG DTYRYI*

black:N5; dark blue:gs linker; cian: nuclear localization sequence; dark green VP16 activation domain; light green: AU1 tag

### P5A: gs linker:NLS:VP16:$^{AU1}$

M ENAALEAKIAALKAKNAALKAEIAALEA GGSGGGGSGGS DPKKKRKV
APPTDVSLGDELHLDGEDVAMAHADALDDFDLDMLGDGDSPGPGFTPHDSAPYGALDMADFEFEQMFTDALGIDEYGG DTYRYI*

black:P5A; dark blue:gs linker; cian: nuclear localization sequence; dark green VP16 activation domain; light green: AU1 tag

### P5: gs linker:NLS:VP16:$^{AU1}$

M SPEDENAALEEKIAQLKQKNAALKEEIQALEYG GGSGGGGSGGS DPKKKRKV
APPTDVSLGDELHLDGEDVAMAHADALDDFDLDMLGDGDSPGPGFTPHDSAPYGALDMADFEFEQMFTDALGIDEYGG DTYRYI*

black:P5; dark blue:gs linker; cian: nuclear localization sequence; dark green VP16 activation domain; light green: AU1 tag

### P5SN: gs linker:NLS:VP16:$^{AU1}$

M ENSQLEEKISQLKQKNSELKEEIQQLE GGSGGGGSGGS DPKKKRKV
APPTDVSLGDELHLDGEDVAMAHADALDDFDLDMLGDGDSPGPGFTPHDSAPYGALDMADFEFEQMFTDALGIDEYGG DTYRYI*

black:CC; dark blue:gs linker; cian: nuclear localization sequence; dark green VP16 activation domain; light green: AU1 tag

### Myc:N5$^{NLS}$

M EQKLISEEDL GEIAALEAKIAALKAKNAALKAEIAALEAGY DPKKKRKV*

red: Myc tag; Black: N5; cian: nuclear localization sequence

### Myc:P5A$^{NLS}$

M EQKLISEEDL ENAALEAKIAALKAKNAALKAEIAALEA DPKKKRKV*

red: Myc tag; Black: P5A; cian: nuclear localization sequence

### Myc:P5$^{NLS}$

M EQKLISEEDL SPEDENAALEEKIAQLKQKNAALKEEIQALEYG DPKKKRKV*

red: Myc tag; Black: P5; cian: nuclear localization sequence

### Myc:P5SN$^{NLS}$

M EQKLISEEDL ENSQLEEKISQLKQKNSELKEEIQQLE DPKKKRKV*

red: Myc tag; Black: P5SN; cian: nuclear localization sequence

### Myc:N5:KRAB

M EQKLISEEDL GEIAALEAKIAALKAKNAALKAEIAALEAGY DPKKKRKV
PKKKRKVDGGGALSPQHSAVTQGSIIKNKEGMDAKSLTAWSRTLVTFKDVFVDFTREEWKLLDTAQQIVYRNVMLENYKNLVSLGYQLTKPDVILRLEKGEEPWLVEREI
HQETHPDSETAFEIKSSV DTYRYI*

red: Myc tag; Black: N5; cian: nuclear localization sequence; orange: KRAB suppressino domain; light green: AU1 tag

### Myc:P5A:KRAB

M EQKLISEEDL ENAALEAKIAALKAKNAALKAEIAALEA DPKKKRKV
PKKKRKVDGGGALSPQHSAVTQGSIIKNKEGMDAKSLTAWSRTLVTFKDVFVDFTREEWKLLDTAQQIVYRNVMLENYKNLVSLGYQLTKPDVILRLEKGEEPWLVEREI
HQETHPDSETAFEIKSSV DTYRYI*

red: Myc tag; Black: P5A; cian: nuclear localization sequence; orange: KRAB suppressino domain; light green: AU1 tag

### Myc:P5:KRAB

M EQKLISEEDL SPEDENAALEEKIAQLKQKNAALKEEIQALEYG DPKKKRKV
PKKKRKVDGGGALSPQHSAVTQGSIIKNKEGMDAKSLTAWSRTLVTFKDVFVDFTREEWKLLDTAQQIVYRNVMLENYKNLVSLGYQLTKPDVILRLEKGEEPWLVEREI
HQETHPDSETAFEIKSSV DTYRYI*

red: Myc tag; Black: P5; cian: nuclear localization sequence; orange: KRAB suppressino domain; light green: AU1 tag

### Myc:P5SN:KRAB

M EQKLISEEDL ENSQLEEKISQLKQKNSELKEEIQQLE DPKKKRKV
PKKKRKVDGGGALSPQHSAVTQGSIIKNKEGMDAKSLTAWSRTLVTFKDVFVDFTREEWKLLDTAQQIVYRNVMLENYKNLVSLGYQLTKPDVILRLEKGEEPWLVEREI
HQETHPDSETAFEIKSSV DTYRYI*

red: Myc tag; Black: P5SN; cian: nuclear localization sequence; orange: KRAB suppressino domain; light green: AU1 tag

### His:TALE-A:NLS:gs lineker:N6 (TALE-A:N6)

M HHHHHH
DYKDHDGDYKDHDIDYKDDDDKMAPKKKRKVGIHRGVPMVDLRTLGYSQQQQEKIKPKVRSTVAQHHEALVGHGFTHAHIVALSQHPAALGTVAVKYQDMIAALPEATHE
AIVGVGKQWSGARALEALLTVAGELRGPPLQLDTGQLLKIAKRGGVTAVEAVHAWRNALTGAPLNLTPDQVVAIASNGGGKQALETVQRLLPVLCQDHGLTPEQVVAIAS
NGGGKQALETVQRLLPVLCQAHGLTPDQVVAIASNIGGKQALETVQRLLPVLCQAHGLTPAQVVAIASHDGGKQALETVQRLLPVLCQDHGLTPDQVVAIASNGGGKQAL
ETVQRLLPVLCQDHGLTPEQVVAIANNNGGKQALETVQRLLPVLCQAHGLTPDQVVAIASHDGGKQALETVQRLLPVLCQAHGLTPAQVVAIASNGGGKQALETVQRLLP
VLCQDHGLTPDQVVAIANNNGGKQALETVQRLLPVLCQDHGLTPEQVVAIASHDGGKQALETVQRLLPVLCQAHGLTPDQVVAIASNGGGKQALETVQRLLPVLCQAHGL
TPAQVVAIASHDGGKQALETVQRLLPVLCQDHGLTPDQVVAIASHDGGKQALETVQRLLPVLCQDHGLTPEQVVAIASHDGGKQALETVQRLLPVLCQAHGLTPDQVVAI
ANNNGGKQALETVQRLLPVLCQAHGLTPAQVVAIASHDGGKQALETVQRLLPVLCQDHGLTPEQVVAIASNGGGRPALESIVAQLSRPDPALAALTNDHLVALACLGGRP
ALDAVKKGLPHAPALIKRTNRRIPERTSHRVAGS DPKKKRKV GGSGGGSGGS KIAALKAEIAALEAENAALEAKIAALKAG*

magenta: his tag; orange: TALA; dark blue: gs linker; cian: nuclear localization sequence; black: N6

### His:TALE-A:NLS:gs lineker:N6:linker:N8 (TALE-A:N6:N8)

M HHHHHH
DYKDHDGDYKDHDIDYKDDDDKMAPKKKRKVGIHRGVPMVDLRTLGYSQQQQEKIKPKVRSTVAQHHEALVGHGFTHAHIVALSQHPAALGTVAVKYQDMIAALPEATHE
AIVGVGKQWSGARALEALLTVAGELRGPPLQLDTGQLLKIAKRGGVTAVEAVHAWRNALTGAPLNLTPDQVVAIASNGGGKQALETVQRLLPVLCQDHGLTPEQVVAIAS
NGGGKQALETVQRLLPVLCQAHGLTPDQVVAIASNIGGKQALETVQRLLPVLCQAHGLTPAQVVAIASHDGGKQALETVQRLLPVLCQDHGLTPDQVVAIASNGGGKQAL
ETVQRLLPVLCQDHGLTPEQVVAIANNNGGKQALETVQRLLPVLCQAHGLTPDQVVAIASHDGGKQALETVQRLLPVLCQAHGLTPAQVVAIASNGGGKQALETVQRLLP
VLCQDHGLTPDQVVAIANNNGGKQALETVQRLLPVLCQDHGLTPEQVVAIASHDGGKQALETVQRLLPVLCQAHGLTPDQVVAIASNGGGKQALETVQRLLPVLCQAHGL
TPAQVVAIASHDGGKQALETVQRLLPVLCQDHGLTPDQVVAIASHDGGKQALETVQRLLPVLCQDHGLTPEQVVAIASHDGGKQALETVQRLLPVLCQAHGLTPDQVVAI
ANNNGGKQALETVQRLLPVLCQAHGLTPAQVVAIASHDGGKQALETVQRLLPVLCQDHGLTPEQVVAIASNGGGRPALESIVAQLSRPDPALAALTNDHLVALACLGGRP
ALDAVKKGLPHAPALIKRTNRRIPERTSHRVAGS DPKKKRKV GGSGGGSGGS KIAALKAEIAALEAENAALEAKIAALKAG GGSGGGSGGS
YKIAALKAENAALEAKIAALKAEIAALEAGC*

magenta: his tag; orange: TALA; dark blue: gs linker; cian: nuclear localization sequence; black: N6 and N8

### CoV-2 Spike-protein

M
FLLTTKRTMFVFLVLLPLVSSQCVNLTTRTQLPPAYTNSFTRGVYYPDKVFRSSVLHSTQDLFLPFFSNVTWFHAIHVSGTNGTKRFDNPVLPFNDGVYFASTEKSNIIR
GWIFGTTLDSKTQSLLIVNNATNVVIKVCEFQFCNDPFLGVYYHKNNKSWMESEFRVYSSANNCTFEYVSQPFLMDLEGKQGNFKNLREVFKNIDGYFKIYSKHTPINLV
RDLPQGFSALEPLVDLPIGINITRFQTLLALHRSYLTPGDSSSGWTAGAAAYYVGYLQPRTFLLKYNENGTITDAVDCALDPLSETKCTLKSFTVEKGIYQTSNFRVQPT
ESIVRFPNITNLCPFGEVFNATRFASVYAWNRKRISNCVADYSVLYNSASFSTFKCYGVSPTKLNDLCFTNVYADSFVIRGDEVRQIAPGQTGKIADYNYKLPDDFTGCV
IAWNSNNLDSKVGGNYNYLYRLFRKSNLKPFERDISTEIYQAGSTPCNGVEGFNCYFPLQSYGFQPTNGVGYQPYRVVVLSFELLHAPATVCGPKKSTNLVKNKCVNFNF

NGLTGTGVLTESNKKFLPFQQFGRDIADTTDAVRDPQTLEILDITPCSFGGVSVITPGTNTSNQVAVLYQDVNCTEVPVAIHADQLTPTWRVYSTGSNVFQTRAGCLIGA
EHVNNSYECDIPIGA
GICASYQTQTNSPRRARSVASQSIIAYTMSLGAENSVAYSNNSIAIPTNFTISVTTEILPVSMTKTSVDCTMYICGDSTECSNLLLQYGSFCTQLNRALTGIAVEQDKNT
QEVFAQVKQIYKTPPIKDFGGFNFSQILPDPSKPSKRSFIEDLLFNKVTLADAGFIKQYGDCLGDIAARDLICAQKFNGLTVLPPLLTDEMIAQYTSALLAGTITSGWTF
GAGAALQIPFAMQMAYRFNGIGVTQNVLYENQKLIANQFNSAIGKIQDSLSSTASALGKLQDVVNQNAQALNTLVKQLSSNFGAISSVLNDILSRLDKVEAEVQIDRLIT
GRLQSLQTYVTQQLIRAAEIRASANLAATKMSECVLGQSKRVDFCGKGYHLMSFPQSAPHGVVFLHVTYVPAQEKNFTTAPAICHDGKAHFPREGVFVSNGTHWFVTQRN
FYEPQIITTDNTFVSGNCDVVIGIVNNTVYDPLQPELDSFKEELDKYFKNHTSPDVDLGDISGINASVVNIQKEIDRLNEVAKNLNESLIDLQELGKYEQYIKWPWYIWL
GFIAGLIAIVMVTIMLCCMTSCCSCLKGCCSCGSCCKFDEDDSEPVLKGVKLHYT*

Dark red:S1; blue: S2

### hACE2

MSSSSWLLLSLVAVTAAQSTIEEQAKTFLDKFNHEAEDLFYQSSLASWNYNTNITEENVQNMNNAGDKWSAFLKEQSTLAQMYPLQEIQNLTVKLQLQALQQNGSSVLSE
DKSKRLNTILNTMSTIYSTGKVCNPDNPQECLLLEPGLNEIMANSLDYNERLWAWESWRSEVGKQLRPLYEEYVVLKNEMARANHYEDYGDYWRGDYEVNGVDGYDYSRG
QLIEDVEHTFEEIKPLYEHLHAYVRAKLMNAYPSYISPIGCLPAHLLGDMWGRFWTNLYSLTVPFGQKPNIDVTDAMVDQAWDAQRIFKEAEKFFVSVGLPNMTQGFWEN
SMLTDPGNVQKAVCHPTAWDLGKGDFRILMCTKVTMDDFLTAHHEMGHIQYDMAYAAQPFLLRNGANEGFHEAVGEIMSLSAATPKHLKSIGLLSPDFQEDNETEINFLL
KQALTIVGTLPFTYMLEKWRWMVFKGEIPKDQWMKKWWEMKREIVGVVEPVPHDETYCDPASLFHVSNDYSFIRYYTRTLYQFQFQEALCQAAKHEGPLHKCDISNSTEA
GQKLFNMLRLGKSEPWTLALENVVGAKNMNVRPLLNYFEPLFTWLKDQNKNSFVGWSTDWSPYADQSIKVRISLKSALGDKAYEWNDNEMYLFRSSVAYAMRQYFLKVKN
QMILFGEEDVRVANLKPRISFNFFVTAPKNVSDIIPRTEVEKAIRMSRSRINDAFRLNDNSLEFLGIQPTLGPPNQPPVSIWLIVFGVVMGVIVVGIVILIFTGIRDRKK
KNKARSGENPYASIDISKGENNPGFQNTDDVQTSF*

### hTMPRSS2 (SinoBiological)

MALNSGSPPAIGPYYENHGYQPENPYPAQPTVVPTVYEVHPAQYYPSPVPQYAPRVLTQASNPVVCTQPKSPSGTVCTSKTKKALCITLTLGTFLVGAALAAGLLWKFMG
SKCSNSGIECDSSGTCINPSNWCDGVSHCPGGEDENRCVRLYGPNFILQVYSSQRKSWHPVCQDDWNENYGRAACRDMGYKNNFYSSQGIVDDSGSTSFMKLNTSAGNVD
IYKKLYHSDACSSKAVVSLRCIACGVNLNSSRQSRIVGGESALPGAWPWQVSLHVQNVHVCGGSIITPEWIVTAAHCVEKPLNNPWHWTAFAGILRQSFMFYGAGYQVEK
VISHPNYDSKTKNNDIALMKLQKPLTFNDLVKPVCLPNPGMMLQPEQLCWISGWGATEEKGKTSEVLNAAKVLLIETQRCNSRYVYDNLITPAMICAGFLQGNVDSCQGD
SGGPLVTSKNNIWWLIGDTSWGSGCAKAYRPGVYGNVMVFTDWIYRQMRADG GGGS EQKLISEEDL*

Black: hMPRSS2; dark blue: gs linker; red:Myc tag

### $GFP_{1\text{-}10}$:gs linker:N7 ($GFP_{1\text{-}10}$:N7)

SKGEELFTGVVPILVELDGDVNGHKFSVRGEGEGDATIGKLTLKFICTTGKLPVPWPTLVTTLTYGVQCFSRYPDHMKRHDFFKSAMPEGYVQERTISFKDDGKYKTRAV
VKFEGDTLVNRIELKGTDFKEDGNILGHKLEYNFNSHNVYITADKQKNGIKANFTVRHNVEDGSVQLADHYQQNTPIGDGPVLLPDNHYLSTQTVLSKDPNEKSGSGSGS
GSSGGSGSGEIAALEAKNAALKAEIAALEAKIAALKAGY*

Green: GFP(1-10); dark blue: gs Linker; black:N7

### $GFP_{1\text{-}10}$:gs linker:P7 ($GFP_{1\text{-}10}$:P7)

SKGEELFTGVVPILVELDGDVNGHKFSVRGEGEGDATIGKLTLKFICTTGKLPVPWPTLVTTLTYGVQCFSRYPDHMKRHDFFKSAMPEGYVQERTISFKDDGKYKTRAV
VKFEGDTLVNRIELKGTDFKEDGNILGHKLEYNFNSHNVYITADKQKNGIKANFTVRHNVEDGSVQLADHYQQNTPIGDGPVLLPDNHYLSTQTVLSKDPNEKSGSGSGS
GSSGGSGSGEIQALEEKNAQLKQEIAALEEKNQALKYG*

Green: GFP(1-10); dark blue: gs Linker; black:P7

### $^{His}$N8:gs linker:$GFP_{11}$ (N8:$GFP_{11}$)

M HHHHHH YGKIAALKAENAALEAKIAALKAEIAALEAGY GGSGGGSG RDHMVLHEYVNAAGIT

magenta: his tag; dark blue: gs linker; black: N8; green: GFP11;

### $^{His}$P8:gs linker:$GFP_{11}$ (P8:$GFP_{11}$)

M HHHHHH KIAQLKE ENQQLEQ KIQALKE ENAALEY GGSGGGSG RDHMVLHEYVNAAGIT

magenta: his tag; dark blue: gs linker; black: P8; green: GFP11

### $^{His}$(N8:gs linker:$GFP_{11}$)$_3$ (3×(N8:$GFP_{11}$))

M HHHHHH YGKIAALKAENAALEAKIAALKAEIAALEAGY GGSGGGSG RDHMVLHEYVNAAGIT GSGGGSG YGKIAALKAENAALEAKIAALKA EIAALEAGY
GGSGGGSG RDHMVLHEYVNAAGIT GGSGGGSG YGKIAALKAENAALEAKIAALKAEIAALEAGY GGSGGGSG RDHMVLHEYVNAAGIT

magenta: his tag; dark blue: gs linker; black: N8; green: GFP11
